# A NOVEL MITOCHONDRIAL PEPTIDE ESSENTIAL FOR RESPIRATORY CAPACITY PROMOTES GROWTH, YIELD AND ABIOTIC STRESS TOLERANCE IN PLANTS

**DOI:** 10.64898/2026.04.13.718140

**Authors:** Lening Gan, Yanqiao Zhu, Xuelu Wang, James Whelan, Huixia Shou

## Abstract

Efforts to boost crop productivity have focused on improving photosynthesis. However, plant respiration consumes over 90% of fixed carbon, fueling growth, stress responses, and resource acquisition. Despite its metabolic centrality, respiratory regulation remains underexplored as a yield target.

Here, we identify AtMLSP1 (At4g17085), a 57-amino-acid mitochondrial inner membrane peptide in Arabidopsis thaliana. Loss-of-function *atmlsp1* mutants show impaired growth, while AtMLSP1 overexpression boosts biomass, seed yield, and high light tolerance.

No induction of alternative oxidase protein abundance or activity was observed in *atmlsp1* plants, a response normally triggered by respiratory perturbation. Mitochondria isolated from *atmlsp1* lines had an approximately 90% reduction in the amount of the intermembrane space protein Cytochrome c and a 30% reduction in all subunits of complex III, while other respiratory complexes remain unaffected. As expected, loss-of-function plants *atmlsp1* reveal a significant decrease in total respiratory capacity and membrane potential, but mitochondrial integrity, and abundance of complex I, II and complex IV were unchanged.

Overexpression of the orthologues of this gene in *Oryza sativa* (rice) and *Glycine max* (soybean) results in a significant increases in yield (seed number and size) under field conditions across different locations.

## INTRODUCTION

A variety of approaches are being undertaken to improve plant productivity in terms of yield and abiotic stress tolerance to meet the goal of global food security. These efforts have largely focused on the improving photosynthetic efficiency at a number of different steps (Croce *et al*., 2024), such as increasing the efficiency of the carboxylation of ribulose bis-phosphate carboxylase oxygenase (Rubisco) (Gionfriddo *et al*., 2024), reducing or by-passing photorespiration (Meacham-Hensold *et al*., 2024), concentrating CO_2_ at the site of fixation by translating algae CO_2_ concentrating mechanisms or engineering C4 photosynthetic pathways into C3 crop plants (Long *et al*., 2018; Furbank *et al*., 2023). However, respiration consumes a significant portion of fixed carbon, along with other energy requiring cost in plants, such as nutrient and water acquisition, transport and associated cost of protein synthesis and degradation, accounting for 90% or more of carbon expenditure (Garcia *et al*., 2023). While modulating respiration is an attractive strategy to potentially increase plant productivity, to date there are no examples where alteration of respiration *per se* has led to increased yield, in terms of seed mass or quantity (Huang *et al*., 2019; Meyer *et al*., 2019; Ghifari *et al*., 2023; Yang *et al*., 2024). Counter-intuitively, over-expression of the Alternative oxidase (AOX) in *Oryza sativa* can increases yield (Nguyen *et al*., 2025).

In fact, there are many cases where AOX has been shown to be beneficial for plant growth, especially under limiting of adverse conditions (Selinski *et al*., 2018), suggesting that growth and stress tolerance is not just a simple process where mitochondria are require to produce ATP via oxidative phosphorylation, and that other factors related to mitochondria may have profound impacts on growth, yield and stress tolerance. The growth versus defense dilemma is well established in plant biology and has traditionally been ascribed to a resource partitioning issue, i.e. energy and metabolites can be used to support either growth or defense processes (Tan *et al*., 2002; He *et al*., 2022). A global analysis of plant resistance genes, commonly referred to as R-genes that are essential for innate immunity compared to specific leaf area as a growth supported this hypothesis for wild type species but not agricultural species (Giolai & Laine, 2024). With reference to activation of mitochondrial retrograde signaling by genetic or pharmacological inhibition of the electron transport chain results in activation of mitochondrial retrograde signaling (MRS), that retards growth, at least in part by alteration of auxin signaling (Ivanova *et al*., 2014; Kerchev *et al*., 2014). Likewise overexpression of the MRS regulating transcription factors ANAC017 retards growth, while its knock-out results in enhanced growth (Ivanova *et al*., 2014). Thus, perturbation of mitochondrial may impair growth through incompatible signaling pathways, rather than resource limitation *per se*.

Small molecular weight proteins and peptides with molecular weights or amino less than 10 kDa or ≤100 amino acids are widely present in eukaryotes. Mitochondrial microproteins in animals play a critical role in regulating energy metabolism. Studies have revealed multiple roles as metabolic sensors, signalling hubs, and functional regulators. SMIM26 is one of the key mitochondrial microproteins localized to the mitochondrial inner membrane and regulates mitochondrial protein translation in response to serine levels, thereby affecting oxidative metabolism (Nah *et al*., 2025). Humanin, the first micro-peptide discovered to be encoded by mitochondrial DNA, has anti-apoptotic and metabolic protective functions. It can improve insulin sensitivity, and is associated with metabolic diseases such as Alzheimer’s disease and diabetes. SHLP2 maintains mitochondrial function through interaction with Complex I and plays a protective role in Parkinson’s disease and obesity-related insulin resistance (Nashine *et al*., 2018; Kim *et al*., 2024).

Here we show a novel mitochondrial-localised small peptide essential for maintenance of respiratory capacity in plants. Over-expression of this peptide enhances tolerance to a variety of abiotic stresses and increased seed yield. Strikingly, overexpression of its orthologs in soybean and rice also boosted grain production under in field conditions. These findings unveil a conserved regulatory mechanism linking mitochondrial function to stress resilience and productivity, offering a promising biotechnological strategy to simultaneously improve crop yield and environmental adaptability.

## MATERIALS AND METHODS

### Plant materials and growth conditions

#### Arabidopsis thaliana

All *Arabidopsis thaliana* plants used in this study were in the Col-0 ecotype. The *atmlsp1-1* T-DNA insertion mutant (SALK_11106C) was purchased from Arabidopsis Biological Resource Centre. The *atmlsp1-2* was generated via CRISPR/Cas9 gene editing using *pHE12.1-zmpl-tRNA(K29)* vector. *AtMLSP1-OE#1*, *AtMLSP1-OE#2* were generated in the lab by inserting the *AtMLSP1* full length coding sequence into the vector *pCAMBIA1300-35S-3×FLAG* and *p35S:AtMLSP1-GFP; atmlsp1-1* was generated in the lab by inserting the *AtMLSP1* full length coding sequence into the vector *pCAMBIA1300-35S-GFP*. For plate-based experiments, seeds were sterilized and stratified for 48 h at 4°C in the dark. Seeds were sown on Gamborg (B5) medium with 1% (w/v) sucrose and 0.8% (w/v) agar and grown under a 14 h of light/10 h of dark or 10 h of light/14 h of dark photoperiod at 23°C with 120 μmol m^-2^ s^-1^ photosynthetic photon flux density. For soil-based experiments, seeds were stratified in ddH_2_O for 48 h at 4°C in the dark and sown in a vermiculite, perlite, and soil mixture (1:1:3). Plants were grown in climate-controlled chambers with the indicated photoperiod (14 h of light/10 h of dark or 10 h of light/14 h of dark), 65% relative humidity, and 120 μmol m^-2^ s^-1^ photosynthetic photon flux density.

#### Glycine Max - Soybean

For soybean, Williams 82 was used as the recipient for Agrobacterium-mediated soybean transformation for overexpression plants driven by *CaMV35S* promoter with GFP tag and CRISPR/Cas9 gene editing, as well as the wild-type control (Song *et al*., 2013). Soybean seeds were sown in pots of two gallons filled with the matrix media. Soybean plants were grown in a greenhouse with day/night temperatures of 30°C/24°C under LD conditions (14 h light/10 h dark, 800 μmol photons m^-2^ s^-1^).

For the field trials, soybean plants were grown at the experimental station of Zhejiang University in Changxin during 2024 Jun to Oct as well as the ‘winter’ nursery station in Sanya during 2024 Nov to 2025 Feb.

#### Oryza sativa - Rice

For rice, Nipponbare was used for physiological experiments and rice transformation as well as GFP-tagged *CaMV35S*-driven *OsMLSP1-*overexpression transgenic plants and CRISPR/Cas9 knock-out mutant *osmlsp1-1/2/3*. Hydroponic experiments were performed using a modified culture solution containing 1430 μM NH_4_NO_3_, 320 μM NaH_2_PO_4_, 510 μM K_2_SO_4_, 1000 μM CaCl_2_, 1640 μM MgSO_4_, 9 μM MnCl_2_, 0.15 μM CuSO_4_, 0.15 μM ZnSO_4_, 0.08 μM (NH_4_)_6_Mo_7_O_24_, 0.02 μM H_3_BO_3_, 125 μM EDTA-Fe, and 250 mM NaSiO_3_ (pH 5.5). Rice seeds were germinated in tap water with 1% nitric acid for 2 days and then transferred to hydroponic culture. Rice seedlings were grown in a growth room with day/night temperatures of 30°C/24°C and a LD conditions (14 h light/10 h dark, 800 μmol photons m^-2^ s^-1^).

For the field trials, rice plants were grown at the winter nursery station in Sanya during 2024 Dec to 2025 Mar.

### High light treatment

For plate-based high light treatment, seeds were sown on B5 medium and put under normal light (120 photons μmol m^−2^ s^−1^) or high light (400 photons μmol m^−2^ s^−1^) directly, phenotypic analysis and detect of ROS were performed using two-week-old seedlings. For NBT staining, seedlings were incubated with 0.5 mg/mL NBT prepared in 10 mM potassium phosphate buffer (pH 7.8) for 2 h and subsequently treated with 90% ethanol at 65°C for 15 min to remove chlorophyll (Ramel *et al*., 2009). For DAB staining, leaves were incubated in 1 mg/mL DAB solution (pH 3.8), as described previously (Ramel *et al*., 2009); chlorophyll was removed as mentioned for NBT. Measurement of total anthocyanins was conducted according to the description (Li *et al*., 2023). Seedling samples were weighed (100 mg) and incubated overnight in 1 mL of HCl-methanol (the volume ratio of methyl alcohol to HCl was 99:1) under 4°C conditions. Anthocyanin extract was then separated from plant tissue by centrifugation at 13000 rpm min^−1^ for 10 min. The absorbance (OD) of the anthocyanin extract at the wavelength of 530 nm and 600 nm was measured by spectrophotometer. Total anthocyanins were calculated by the formula (A530-A600) g^−1^ FW.

### Phenotypic analysis

Arabidopsis - A Canon digital camera (EOS M50 Mark II) was used to capture images of the whole plants, detached leaves and roots, flowers or siliques or root hair related images were captured by a stereomicroscope (Cqoptec, SZ680), and the root length was measured using Image J.

Soybean - A Canon digital camera mentioned above was used to capture images of the whole plants and seeds, and flowers related images were captured by a stereomicroscope mentioned above.

Rice - A Canon digital camera mentioned above was used to capture images of the whole plants, grains and seeds.

### Construction of vectors

Vectors for transformation of Arabidopsis were constructed as outlined above.

For overexpression constructs used in transformation of soybean, the full length CDSs of *GmMLSP1* and *GmMLSP2* were insert into *pTF101* by seamless cloning. *GFP* was fused at the C terminus with target genes and driven by the *CaMV35S* promoter. The *CRISPR/Cas9* construct was based on the *pBlu-gRNA* vector with gRNA and *CAS9-MDC123* with Cas9 protein. The target sequence (5’- GTTCAGACTATTGCCACTGCTGG-3′) was designed in a region conserved in both *GmMLSP1* and *GmMLSP2* in the first exon of the coding sequence. The target sequence was synthesized and cloned into *pBlu-gRNA* and named *pBlu-gRNA-gmmlsp1/2*. The gRNA cassette with target sequence from *pBlu-gRNA-gmmlsp1/2* was cloned into the *Cas9-MDC123* vector and named *Cas9-gmmlsp1/2*. The constructs were transformed into *Agrobacterium* strain *LBA4404* for soybean transformation.

For rice, the full length CDS of *OsMLSP1* was insert into *pCAMBIA1300* by seamless cloning. *GFP* was fused at the C terminus with target genes and driven by the *CaMV35S* promoter. CRISPR/Cas9 vector for knock out of *OsMLSP1* was constructed by Biogle Genetech (Hangzhou Biogle Co. LTD.) as well as transformation of it and overexpress vector mentioned above.

### Subcellular localization and detection of mitochondrial membrane potential in root of Arabidopsis

For subcellular localization, seven-day-old *p35S:AtMLSP1-GFP; atmlsp1-1* Arabidopsis seedlings were used for co-localization of AtMLSP1-GFP fusion protein and tetramethyl rhodamine methyl ester (MedChemExpress, TMRM). The roots were stained in 1 μM TMRM for stained for 20 min, followed by a single wash with ddH_2_O for 5 min.

For protoplast, transgenic line *p35S:AtMLSP1-GFP; atmlsp1-1* was used for generating protoplasts. Protoplasts were isolated as previously described (Yoo *et al*., 2007). Approximately 10 μg of the COX IV-mCherry was used for each transformation. Protoplasts for confocal microscopy imaging were incubated at 23°C in darkness for 12∼14 h before imaging.

For detection of mitochondrial membrane potential, seven-day-old Col-0, *atmlsp1-1*, *atmlsp1-2*, *AtMLSP1-OE#1* and *AtMLSP1-OE#2* seedlings were stained in both 1 μM TMRM and Mito-Tracker Green FM (Invitrogen) for 20 min, followed by a single wash with ddH_2_O for 5 min (Canal *et al*., 2024).

Confocal fluorescence images were acquired using a LSM 710 NLO confocal laser scanning microscope (Zeiss, Göttingen, Germany), while images of high resolution were acquired with LSM 880. Excitation/emission wavelengths were 488 nm/490–550 nm for GFP and Mito-Tracker Green, 561 nm/575–630 nm for mCherry, 552 nm/590-640 nm for TMRM. Images were processed with Zeiss imaging software (ZEN, blue edition, v.2.3) and Fiji.

### Isolation of mitochondria, gel electrophoresis and immunoblotting

Mitochondria were isolated from 2-week-old seedlings grown on Gamborg (B5) medium plates under LD conditions. Mitochondrial isolation was combined with differential centrifugation and Percoll density gradient centrifugation following a previously published protocol (Lyu *et al*., 2018).

For BN-PAGE, samples were prepared by solubilizing 20 μg mitochondria with 5% (w/v) digitonin as previously described (Eubel *et al*., 2005). Samples were resolved using precast BN-PAGE 4% to 16% Bis-Tris gels (Invitrogen). Coomassie stain, CI and CIII in-gel activity staining were carried out as previously described (Smet *et al*., 2011; Huang *et al*., 2015).

For western blot, 20 μg mitochondria were separated by SDS-PAGE and transferred to a PVDF membrane (0.22 μm, Bio-Rad). Immunodetections were performed as described previously (Li *et al*., 2019). The antibodies used in this study are listed in Supplemental Dataset S3. The intensities of bands were quantified using Image Lab software (Bio-Rad) and calculated relative to the wild type. Three biological replicates were performed.

### Respiratory Measurements

Oxygen consumption of purified mitochondria was measured by a computer-controlled Clark-type O_2_ electrode (Hansatech Instruments, Pentney, UK) according to a previously published protocol (Lyu *et al*., 2018).

All reactions were carried out at 25°C using 950 μL of mitochondrial reaction medium (0.3 M sucrose, 10 mM TES, 10 mM NaCl, 4 mM MgSO_4_, 0.1% [w/v] BSA [pH 7.2]) and mitochondria equivalent to approximately 100 mg of protein. Three biological replicates were performed for each treatment.

### RNA Isolation And Quantitative RT (qRT)-PCR

Total RNA was isolated from plant samples using the RNA-Easy isolation reagent (Vazyme) according to the manufacturer’s recommendations. First-strand cDNAs were synthesized from total RNA using a Primescript RT Reagent Kit with gDNA eraser, and qRT-PCR was performed using TB Green qPCR Master Mix (Takara) on a Lightcycler 480 Thermocycler (Roche Diagnostics). Relative gene expression was calculated by the 2^−ΔΔct^ method using the housekeeping gene *AtActin*, *GmActin* or *OsActin* for different plants as an internal reference. Primers used for qRT-PCR are given in Supplemental Dataset S4. All experiments were performed with three biological And two technical replications.

### Outer Membrane–Ruptured Mitochondria Preparation And Proteinase K Treatment

First, protein concentration of mitochondria should be adjusted to 2 μg/μL, then 1000 μg/500 μL sample was precipitated, and the pellet was gently resuspended in 100 μL of SEH buffer (250 mM sucrose, 1 mM EDTA, and 10 mM HEPES-KOH, 0.1% [w/v] BSA , pH 7.4) and incubated on ice for 15 min after mixing with 1550 μL of 20 mM HEPES-KOH (0.1% [w/v] BSA, pH 7.4). Following incubation, 250 μL of 2 M sucrose and 100 μL of 3 M KCl were added to rupture the outer membrane. Equal amounts of outer membrane-ruptured mitochondria and intact mitochondria were incubated for 30 min on ice with 0 to 1 mg/mL proteinase K, and 2 μL of 100 mM PMSF plus AEBSF was added to terminate the reaction. Samples were precipitated by centrifugation at 13,000 g at 4°C for 3 min and resuspended in 100 μL of SDS-PAGE sample buffer. 5 μL of each sample was used for SDS-PAGE and immunodetection (Schäfer *et al*., 2022).

### Carbonate Extraction

Mitochondrial pellets were resuspended in 0.1 M Na_2_CO_3_ and incubated on ice for 30 min followed by centrifugation of 30 min at 20,000 g at 4°C. The pellet taken as the membrane fraction was resuspended in SDS-PAGE sample buffer and boiled for 5 min at 95°C, while the supernatant was taken as the soluble fraction. The supernatant was transferred into a new 1.5 mL microcentrifuge tube, and 0.1 vol 100% TCA was added to the supernatant to precipitate soluble mitochondrial proteins. After incubating for 15 min on ice, soluble fraction was collected by centrifugation of 15 min at 20,000 g at 4°C, and pellet was resuspended in SDS-PAGE sample buffer and boiled for 5 min at 95°C after washed by ice cold 80% (v/v) acetone for two times and air dried for 5 min (Schäfer *et al*., 2022).

### Statistical Analyses

The data were analysed using one-way ANOVA. At least 3 biological replicates were used for the calculation of SD. Statistical analysis and graphics were carried out with GraphPad Prism software. The statistical analysis for each experiment is described in the Figure legends.

## RESULTS

### AtMLSP1 is a mitochondrial peptide essential for respiratory chain function

We identify a novel 57-amino acids peptide (encoded by *At4g17085*) as a critical regulator of mitochondrial respiratory capacity. Previously five independent mass spectrometry analyses with purified mitochondria confirm its mitochondrial localisation (Hooper *et al*., 2017), and ComplexomeMap identifies it on blue native polyacrylamide gel electrophoresis (BN-PAGE) gels (Senkler *et al*., 2017) (Fig. S1). We confirmed the subcellular localization of this protein using both a targeting and accumulation approaches (Millar *et al*., 2009). GFP tagging visualisation using Arabidopsis root hairs and protoplast displayed a clear overlap with the mitochondrial marker TMRM and COXIV-mCherry respectively (Fig. 1a,b), supporting the independent mass spectrometry studies. To confirm a mitochondrial localisation, we used immunoblotting with anti-Flag and anti-GFP antibodies of tagged lines on purified mitochondria. Both displayed a clear distinct signal showing that the protein encoded by *At4g17085* could target and accumulate in mitochondria (Fig. 1c). Thus, we called this peptide Mitochondrial Localised Small Peptide 1 (AtMLSP1). To determine the intraorganellar location we utilised a combination of carbonate extraction and protease accessibility assays (Fig. 1d,e). Carbonate extraction revealed a clear membrane location with the protein being extracted and located in the pellet fraction (Fig. 1d). Protease accessibility assays used Tom20-3 as an outer membrane marker, Cytochrome c as an intermembrane space marker, Tim23-1 as an inner membrane marker and AOX as a matrix marker (an interfacial membrane protein located on the matrix side of the inner membrane). Both Flag and GFP tagged MLSP1 were only digested at the highest level of protease, supporting an inner membrane protected location.

**Figure 1.**
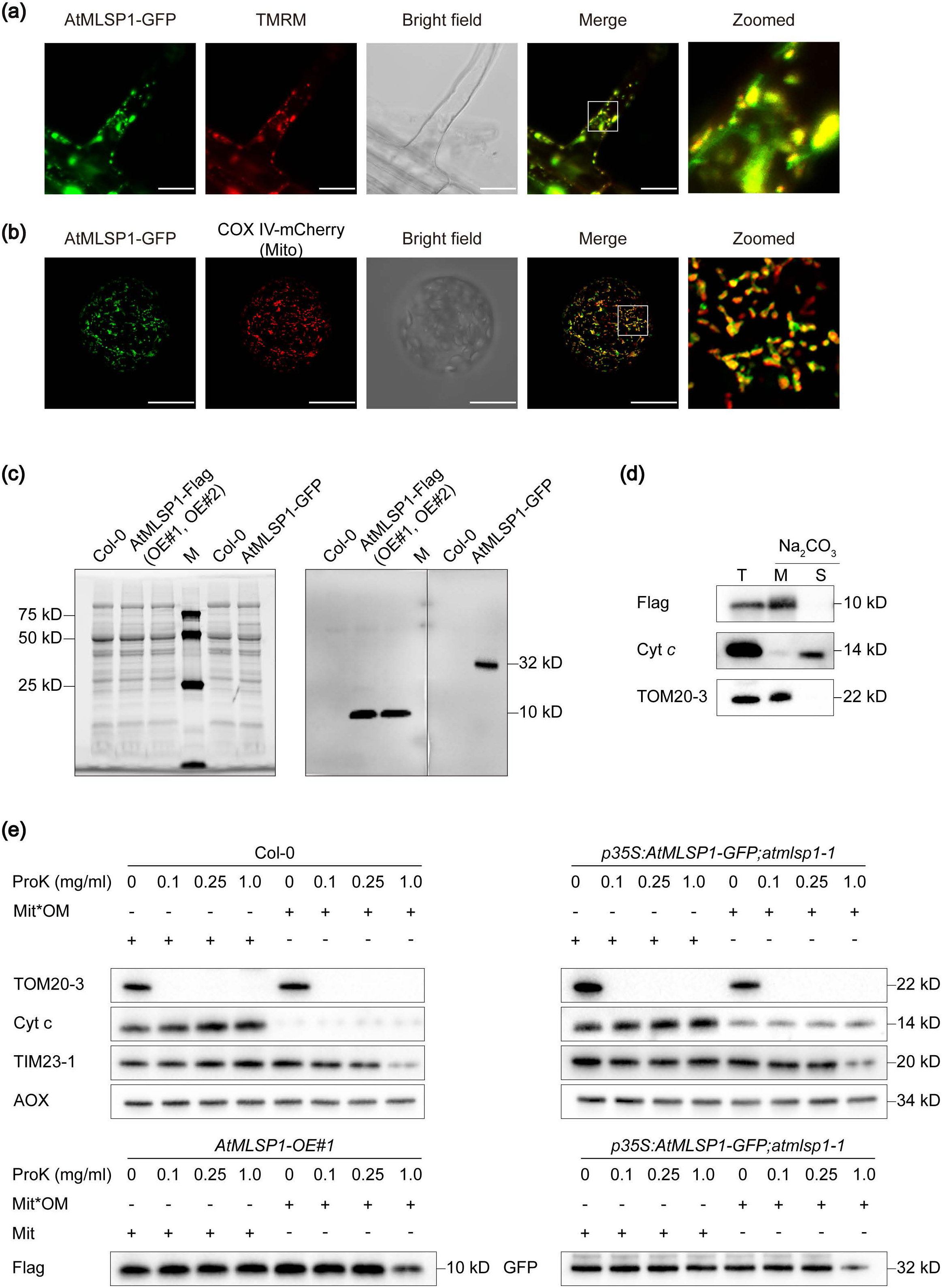
AtMLSP1 is a mitochondrial inner membrane protein required to maintain complex III (cytochrome bc_1_) and Cytochrome c abundance in *Arabidopsis thaliana*. **(a)** Confocal images showing the subcellular localization of AtMLSP1 in Arabidopsis root cells. The *p35S:AtMLSP1-GFP; atmlsp1-1* transgenic seedling was stained with TMRM. Scale bar = 20 µm. **(b)** Confocal images showing the subcellular localization of AtMLSP1 in Arabidopsis protoplast from *p35S:AtMLSP1-GFP; atmlsp1-1* transgenic seedlings. COX IV-mCherry acts as a mitochondrial marker. **(c)** Image of purified mitochondrial proteins of Col-0, *AtMLSP1-OE#1*, *AtMLSP1-OE#2* and *p35S:AtMLSP1-GFP; atmlsp1-1* and immunodetection using anti-FLAG and anti-GFP antibody. **(d)** Carbonate extraction of mitochondria purified from AtMLSP1-OE1 line where MLSP1 is tagged with the Flag epitope. T is total mitochondrial fraction, M is membrane (pellet) fraction that has been solubilised after carbonate extraction and S is soluble fraction that is neutralised. Separation of proteins by SDS-PAGE, blotting and probing with the antibodies indicated. **(e)** Protease accessibility assays on mitochondria purified from wild type (Col-0), a Flag tagged overexpressing line for MLSP1 (At*MLSP1-OE1*) and a knock-out line that has been complemented with a CaMV 35S promoter driving the expression of *AtMLSP1.* ProK = Proteinase K, Mit*OM is outer membrane ruptured mitochondria, Mit is mitochondria. The antibodies used to probe the protein separated by SDS-PAGE and blotted are indicated. TOM20-3 (Translocase of the outer membrane) is an outer membrane protein, Cyt c (Cytochrome c) is an intermembrane space protein, TIM23-1 (Translocase of the inner membrane) is an inner membrane protein and AOX (Alternative oxidase) is an interfacial membrane protein on the inside of the inner membrane).

### AtMLSP1 maintains respiratory supercomplex stability

We analysed the impact of knocking-out and overexpressing *MLSP1* using BN-PAGE analysis and western blot analysis. BN-PAGE of the stained respiratory complexes revealed a marked reduction in the stained band intensities in dimeric complex III and supercomplex of I and III_2_ bands in the two independent knock-out lines, and showed no much difference between Col-0 and overexpress lines (Fig. 2a, left panel – marked with asterisk). Activity staining for complex III confirmed the reduction in dimeric complex III and supercomplex I+III_2_ in two mutant lines (Fig. 2a, middle panel – marked with asterisk). Activity stain of complex I and quantification of those bands were also performed, two mutant lines showed a decrease in intensity of supercomplex I+III_2_, in agreement with the staining of complex III, with complex I was almost unaffected, but noticeably supercomplex I+III_2_+IV had increased, while activity of supercomplex I+III_2_ as well as I+III_2_+IV showed significant increase in two overexpress lines (Fig. 2a right panel – asterisk mark for supercomplex I+III, black dot mark for complex I and supercomplex I+III_2_+IV and Fig.2b). Western blot analysis of purified mitochondria purified from wild type, two independent *atmlsp1* lines and At*MLSP1-OE* lines confirmed the changes observed with BN-PAGE (Fig. 2c). Probing mitochondria with NDUFS4 (complex I), SDH1 (complex II), RISP (complex III) COXII (complex IV) and ATPb (complex V), as well as Cyt *c*, AOX and TOM40, the latter as a loading control, revealed that the abundance of cytochrome c was reduced by almost 90% and RISP that detected the Rieske FeS protein of complex III by ∼30%. Notably, AOX was not greatly increased in abundance.

**Figure 2.**
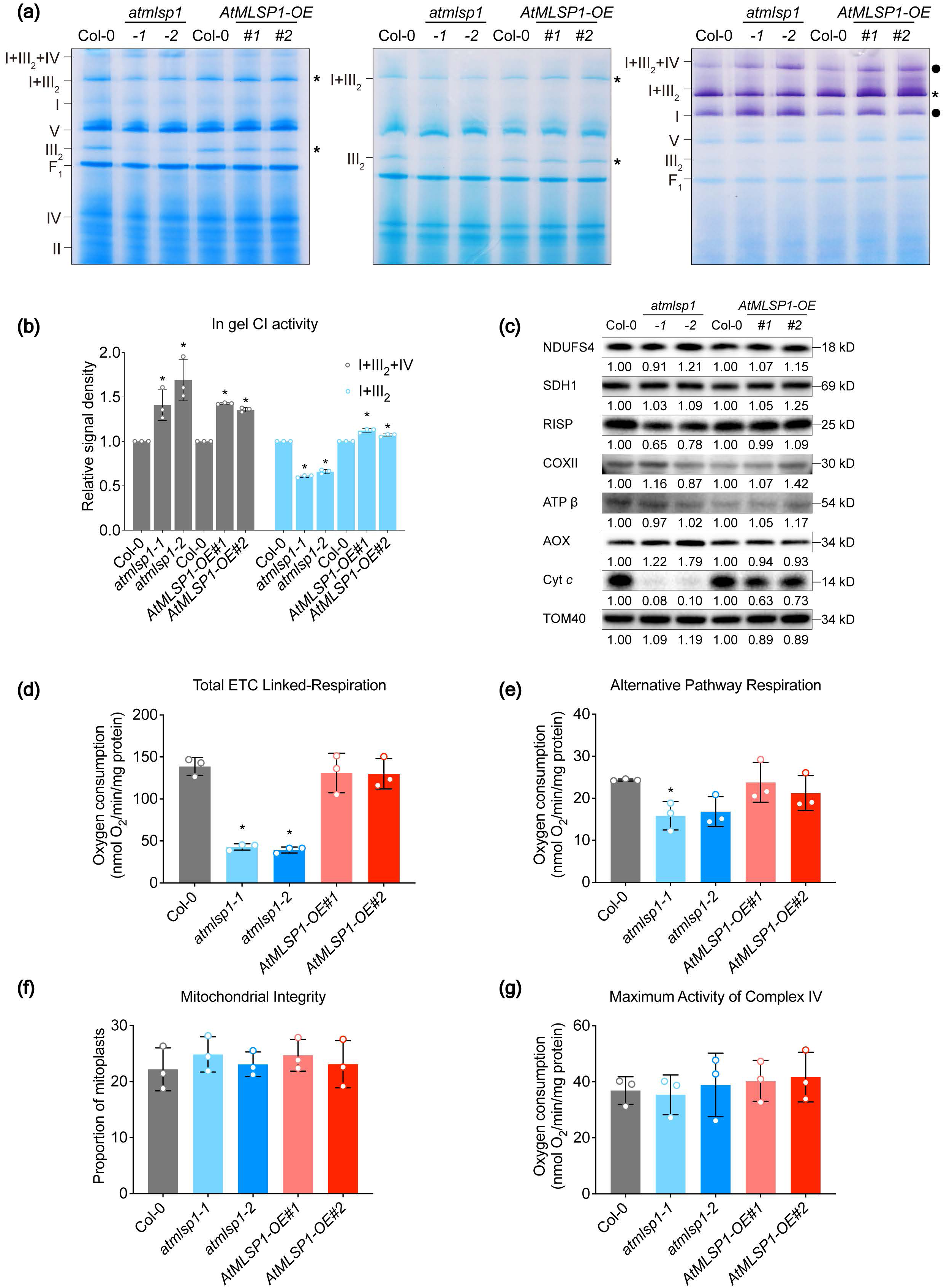
Biochemical characterisation of mitochondria purified from loss-of-function *atmlsp1* and overexpression AtMLSP1 lines. **(a)** Mitochondria (20 µg) were isolated from all lines and resolved by BN-PAGE and stained with Coomassie blue (left panel), in-gel activity for complex III (middle panel) and complex I (right panel). **(b)** As the intensity of the staining of complex I activity appears to vary from supercomplex I + III_2_ and I+III_2_+IV between the various mutant lines and Col-0, the staining intensity of each band was quantified. Data were expressed as means ± SD from three biologicals. Significant difference (*P < 0.05) based on one-way ANOVA compared with Col-0 are indicated by asterisks. **(c)** The protein abundance of respiratory chain components from purified mitochondria separated by SDS-PAGE, blotted and probed with antibodies to subunits of each respiratory chain complex. Mitochondrial proteins (20 µg) were loaded. NDUFS4 – complex I, SDH1 – complex II, RISP – complex III, COXII – complex IV, ATPβ – complex V, AOX – Alternative oxidase, Cyt c – Cytochrome c. TOM40 – Translocase of the outer membrane 40 as a loading control. **(d-g)** Oxygen consumption of mitochondria purified measured using a Clark-type oxygen electrode. Total ETC-Linked respiration of mitochondria, purified from Col-0, mutants and overexpressing plants, Alternative pathway respiration of mitochondria, integrity of purified mitochondria purified maximum complex IV activity of purified mitochondria from Col-0, mutants and overexpressing plants. Values are means ± SDs (n = 3; one-way ANOVA; *P < 0.05).

Following the reduced abundance of complex III, cytochrome respiratory capacity was measured to assess the functional consequence. Total oxygen consumption was significantly reduced by approximately 70% in the *atmlsp1* knockout lines, but no significant change between Col-0 and over-expression lines (Fig. 2d). Respiration via the alternative oxidase was also reduced by ∼ 35% (Fig. 2e). Given the severe respiration defect observed, we checked the integrity of mitochondria and cytochrome c oxidase capacity (complex IV) from all the lines. While the respiratory capacity was significantly decreased in the knock-out lines, no differences in the integrity of mitochondria or Cytochrome c oxidase capacity were detected between the lines (Fig. 2f,g). These activities are consistent with the protein abundances observed above. We also monitored the membrane potential using the membrane potential dye TMRM, to determine if there were any differences between the lines. In wild type plants, localization of the two markers was highly consistent (Fig. 3), and while occasionally a mitochondrion could be observed that did not have a corresponding membrane potential. This may be due to the mitoflash phenomena that has been described previously (Demaurex & Schwarzlander, 2016; Booth *et al*., 2021). Analysis of the membrane potential from *atmlsp1* plants revealed that there was a large reduction in the membrane potential, with values only reaching approximately 30% of those of wildtype. When *AtMLSP1* was overexpressed, the mitochondrial membrane potential appeared consistently higher than wildtype (Fig. 3).

**Figure 3.**
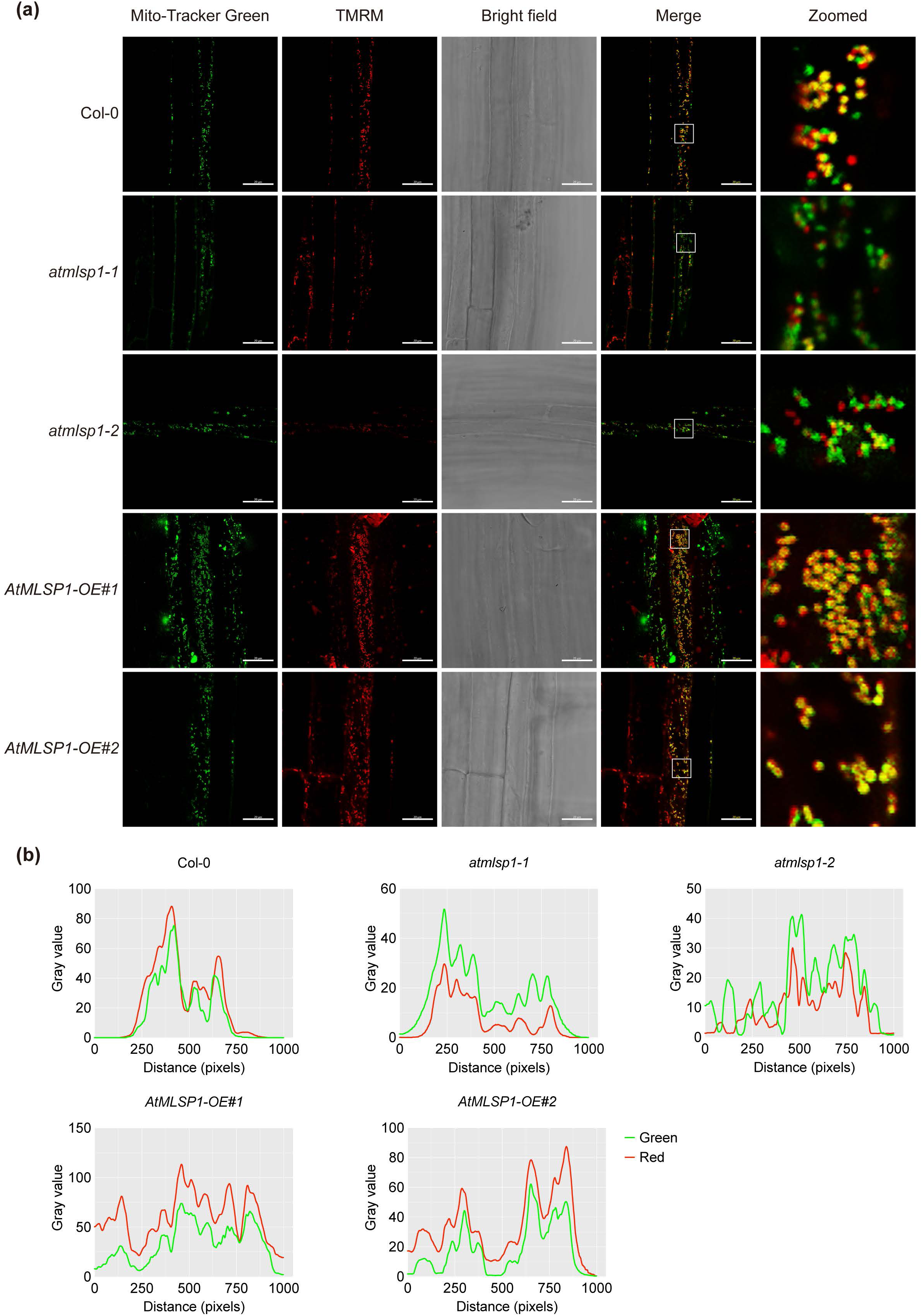
Membrane potential measurement from loss-of-function *atmlsp1* and overexpression AtMLSP1 lines in Arabidopsis roots. **(a)** Detection of mitochondrial membrane potential in roots of Col-0, mutants and overexpressing plants stained with 1 μM Mito-Tracker Green and tetramethyl rhodamine methyl ester (TMRM). Scale bar = 20 μm. **(b)** Gray value of zoomed area in roots of Col-0, mutants and overexpressing plants. Green lines show gray value of Mito-Tracker Green and red lines for TMRM.

The various assays carried out show that AtMLSP1, BN-PAGE abundance and activity assays, western blot analysis and respiratory measurements all show that AtMLSP1 is required to maintain respiratory competence in mitochondria. A loss-of-function mutant results in a large decrease in cytochrome supported respiration, but AOX is not induced.

### Over-expression of *AtMLSP1* enhances growth, seed yield and stress tolerance

Mutation of genes encoding components of the mitochondrial respiratory chain generally results in retarded growth or even lethality at a gametophyte or embryonic stage (Welchen *et al*., 2012; Zhu *et al*., 2012; Dahan *et al*., 2014; Kuhn *et al*., 2015). Analysis of knock-out mutants for *AtMLSP1* displayed a retarded growth phenotype, at all stages of growth, with fewer leaves and reduced seed yield (Fig.4a). Growth of over-expression lines didn’t seem to be inhibited, but differed in some aspects compared to Col-0 plants (Fig.4a). Growth in terms of leaf emergence for approximately the first 40 days of growth was slightly retarded (Fig. S2a). However, after approximately 40 days, the overexpressing lines continue to growth so that after not only had gained on Col-0 plants in terms of overall size (Fig. 4a) but also produced more leaves (Fig. 4a and Fig. S2a). To examine this more closely we analyzed growth on agar plates with stratified seeds to examine where growth was retarded (Fig. S2b and c). While the loss-of-function line for *atmlsp1* was clearly delayed from radical emergence, the difference between wild type and *AtMLSP1-OE* lines was more subtle, so that leaf emergence was delayed 1 to 2 days behind wildtype, so that at stage 1.10, *AtMLSP1-OE* was still ∼ 1 days developmentally delayed compared to wild type, while *atmlsp1* was ∼ 7 days. Also *AtMLSP1-OE* lines continue to grow and after wildtype had transitioned to flowering, and by day 64 was larger in both vegetive mass, and notably seed yield was significantly increased (Fig. 4a).

**Figure 4.**
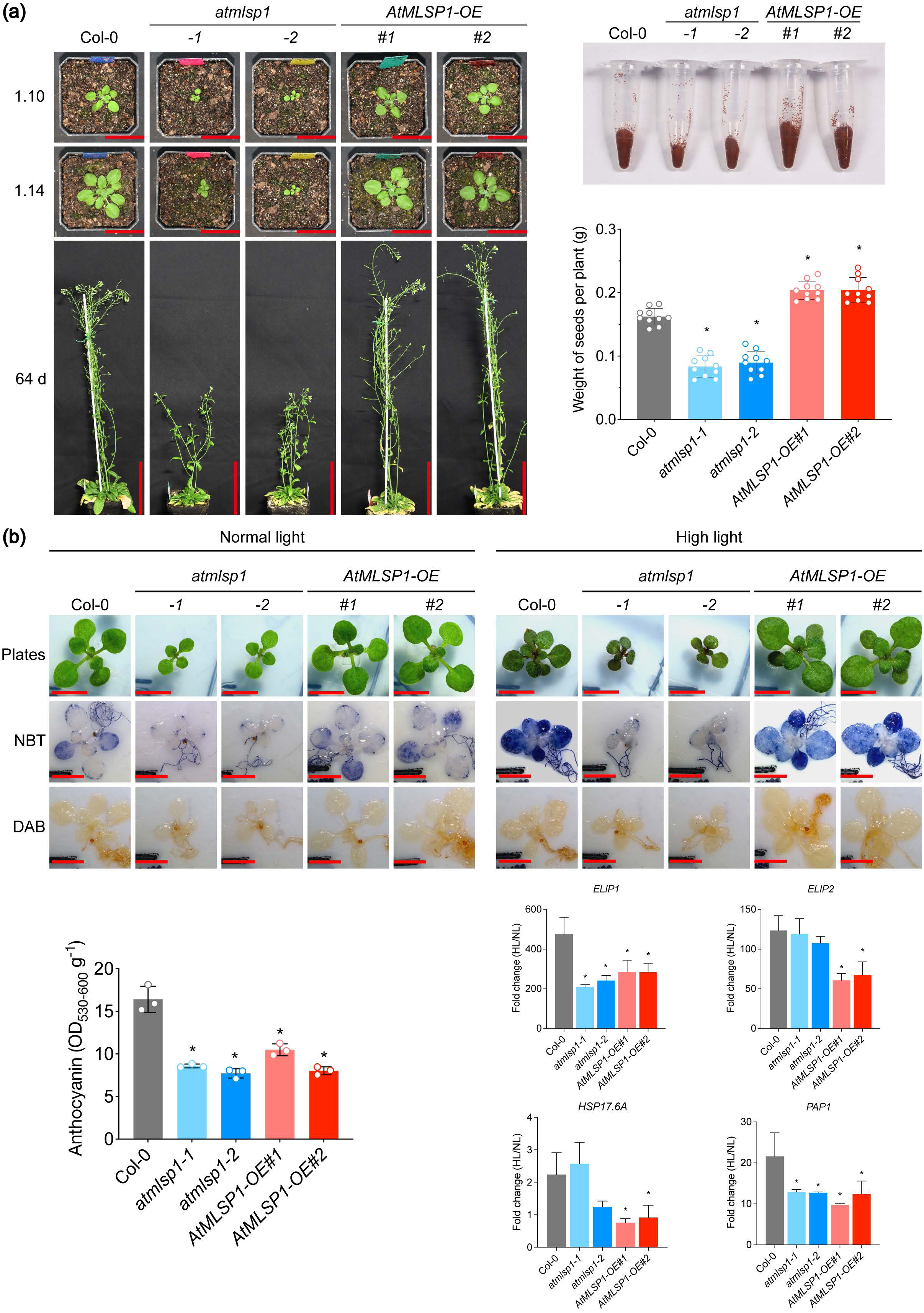
Phenotypic analysis *Arabidopsis thaliana* loss-of-function and overexpressing plants. **(a)** Representative images from the stage growth progression analysis from Col-0, mutants and overexpressing plants. 1.10 (10 rosette leaves > 1 mm in length), 1.14 (14 rosette leaves > 1 mm in length). Scale bar, 3 cm for 1.10 and 1.14 stage, and 10 cm for 64 d. Seed yield of Col-0, mutants and overexpressing plants are shown. **(b)** Phenotype of Col-0, mutants and overexpressing seedlings grown in plates under control treatment and high light stress at 2 weeks. Scale bar, 5 mm. Staining with NBT for Superoxide ions (O^2-^) and DAB for hydrogen peroxide (H_2_O_2_) is shown. Scale bar = 1 mm. Anthocyanin accumulation in rosette leaves of Col-0, mutants and overexpressing plants after 9 d treatment of high light. Fold-changes in transcript abundance for *early light inducible protein 1* and *2* (*ELIP*), *heat shock protein 17.6A* (*HSP 17.6A*), and *production of anthocyanin pigment 1* (*PAP1*). Values are means ± SDs (n = 3; one-way ANOVA; *P < 0.05).

The response of the knockout and overexpression lines to various abiotic high light conditions was examined. Plants were exposed to high light treatment of 400 photons μmol m^−2^ s^−1^ that was almost four times of the permissive growth conditions. Again, greater tolerance was observed with the overexpression lines in terms that they appeared to have less anthocyanins, that was confirmed calorimetrically (Fig. 4b). Notably the knockout lines had reduced anthocyanins as well. To investigate this further experiment was carried out on a plate-based assay and plants stained for ROS with NBT for superoxide ions (O^2-^) and DAB for hydrogen peroxide (H_2_O_2_) after 9 days on high light (Fig. 4b). Intense staining with NBT was observed in Col-0 and was reduced in overexpressing lines, but little staining was observed in the knock-out lines. Thus, the overexpressing lines respond to increased light, but this response is modulated. Using defined markers for high light response, early genes that *encode light inducible protein 1* and *2* (*ELIP*), *heat shock protein 17.6A* (*HSP 17.6A*), and *Production of Anthocyanin Pigment 1* (*PAP1*), markers for light stress it was evident that the response of the mutant lines differed to wild type. For the overexpressing lines *AtMLSP1-OE*, the induction of all genes was significantly reduced compared to wild type, consistent with reduced ROS and anthocyanin production. For *atmlsp1* lines a significant reduction was observed for *ELIP1* and *PAP1,* the latter consistent with the reduction in anthocyanins observed.

Combined these results show that overexpression of *AtMLSP1* results in altered growth and greater tolerance to high light. Furthermore, the altered growth results in increased seed yield. Thus, despite the increase in abiotic stress tolerance, there appears to be no growth or yield drag for overexpression of this protein in plants, prompting investigation of how it would perform in crop plants.

### Over-expression of rice and soybean orthologues resulted in significant increase in seed yield

Since *AtMLSP1* promotes plant growth, we investigated whether modifying its expression in crop species could improve yield. We identified a homologous gene of *AtMLSP1* in rice and two homologs in soybean and generated transgenic materials, including mutants and overexpression lines for both species (Fig. S4 and S5).

Under greenhouse conditions, rice mutant lines exhibited decreased plant height, primary panicle length, seed size, and single-plant yield compared to the wild-type (Nip), whereas overexpression lines showed no significant differences (Fig. 5a, Fig. S6a,b). Strikingly, field-grown rice displayed more pronounced phenotypic effects. Mutants showed shorter primary panicles with increased hollow grains and reduced grain yield, while overexpression lines exhibited longer primary panicles and increased grain yield (Fig. 5b,c, S7 and Table 1). Further, the overexpression lines showed significant increases in plant height, number of effective panicle number, grains per plant, yield per plant, and thousand-grain weight compared to the control, whereas the mutant displayed the opposite trends (Fig. 5b,c, S7 and Table 1).

**Figure 5.**
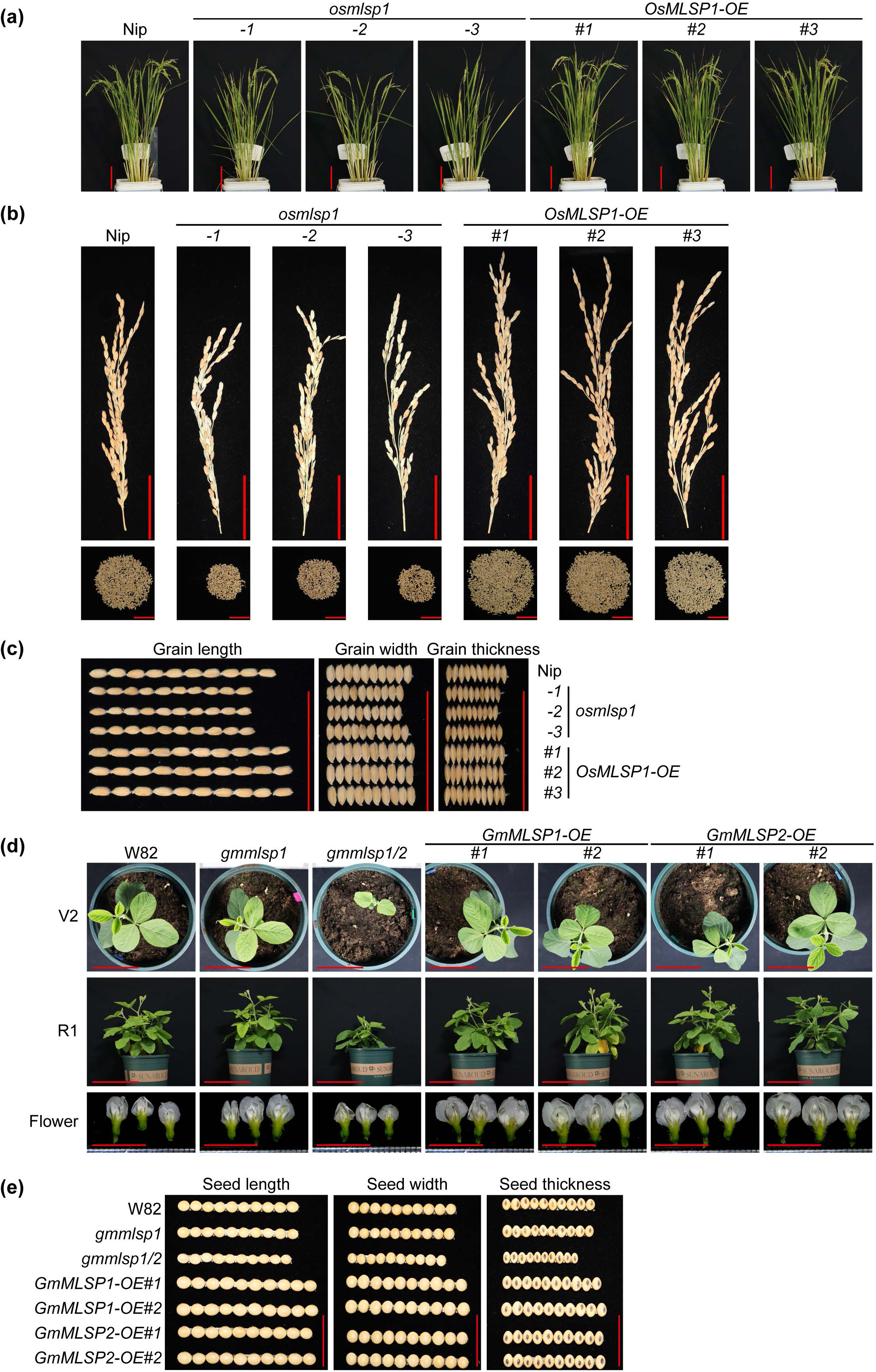
*MLSP* overexpressing transgenic rice and soybean exhibit improved growth and yield. **(a)** Growth phenotypes of Nipponbare, loss-of-function and *MLSP1-*overexpressing rice plants in the green house at 125 d after sowing. Scale bar, 10 cm. **(b)** Primary panicles and seed production per plant of Nipponbare, loss-of-function and *MLSP1-*overexpressing rice. Image displays striking differences in harvested seed numbers from individual plants. Scale bar, 5 cm. **(c)** Seed morphology of Nipponbare, mutants and *MLSP1*-overexpressing rice. Scale bar, 5 cm. **(d)** Vegetative and floral development of W82, mutants and *MS* overexpressing soybean plants under greenhouse conditions. V2, the first fully expanded trifoliate leaf. Scale bar, 11.5 cm; R1, the beginning bloom stage. Scale bar, 23 cm; Flowers photos were taken at R2 (full bloom stage) in green house. Scale bar, 1 cm. **(e)** Seed morphology of W82, mutants and *MLSP* overexpressing soybean plants. Seeds were harvested from the field experiment in Hainan. Scale bar, 3 cm.

**Table 1.**
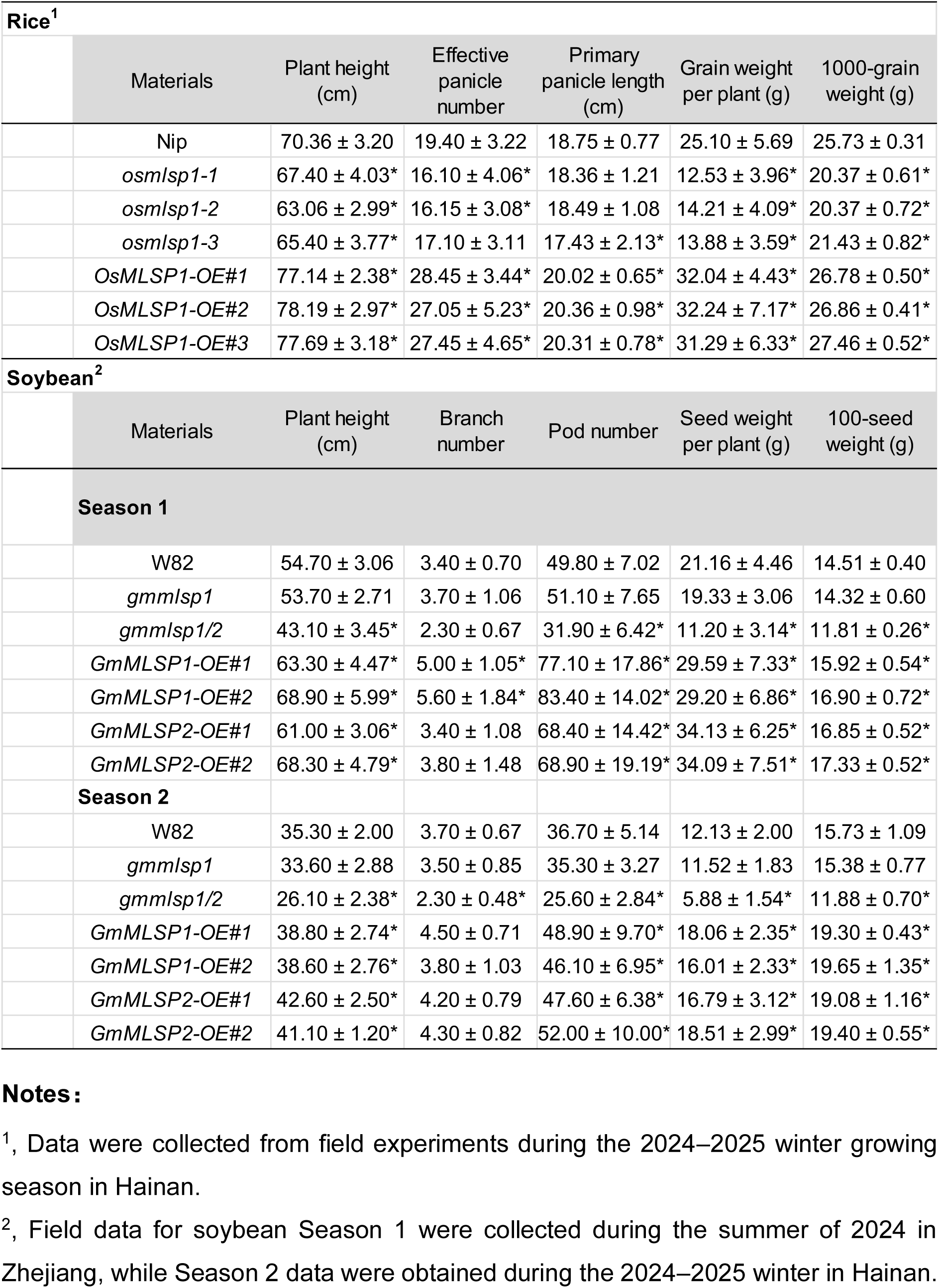
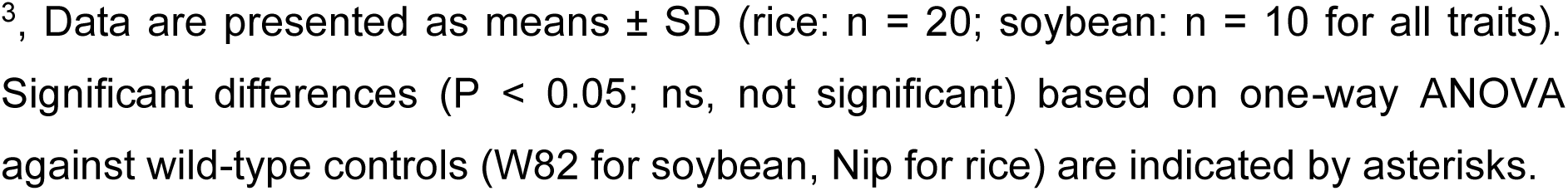
Phenotypic performance of *MLSP*-overexpression rice and soybean.

| Rice<sup>1</sup> |  |  |  |  |  |  |
| --- | --- | --- | --- | --- | --- | --- |
|  | Materials | Plant height (cm) | Effective panicle number | Primary panicle length (cm) | Grain weight per plant (g) | 1000-grain weight (g) |
|  | Nip | 70.36 ± 3.20 | 19.40 ± 3.22 | 18.75 ± 0.77 | 25.10 ± 5.69 | 25.73 ± 0.31 |
|  | *osmlsp1-1* | 67.40 ± 4.03* | 16.10 ± 4.06* | 18.36 ± 1.21 | 12.53 ± 3.96* | 20.37 ± 0.61* |
|  | *osmlsp1-2* | 63.06 ± 2.99* | 16.15 ± 3.08* | 18.49 ± 1.08 | 14.21 ± 4.09* | 20.37 ± 0.72* |
|  | *osmlsp1-3* | 65.40 ± 3.77* | 17.10 ± 3.11 | 17.43 ± 2.13* | 13.88 ± 3.59* | 21.43 ± 0.82* |
|  | *OsMLSP1-OE#1* | 77.14 ± 2.38* | 28.45 ± 3.44* | 20.02 ± 0.65* | 32.04 ± 4.43* | 26.78 ± 0.50* |
|  | *OsMLSP1-OE#2* | 78.19 ± 2.97* | 27.05 ± 5.23* | 20.36 ± 0.98* | 32.24 ± 7.17* | 26.86 ± 0.41* |
|  | *OsMLSP1-OE#3* | 77.69 ± 3.18* | 27.45 ± 4.65* | 20.31 ± 0.78* | 31.29 ± 6.33* | 27.46 ± 0.52* |
| Soybean<sup>2</sup> |  |  |  |  |  |  |
|  | Materials | Plant height (cm) | Branch number | Pod number | Seed weight per plant (g) | 100-seed weight (g) |
| Season 1 |  |  |  |  |  |  |
|  | W82 | 54.70 ± 3.06 | 3.40 ± 0.70 | 49.80 ± 7.02 | 21.16 ± 4.46 | 14.51 ± 0.40 |
|  | *gmmlsp1* | 53.70 ± 2.71 | 3.70 ± 1.06 | 51.10 ± 7.65 | 19.33 ± 3.06 | 14.32 ± 0.60 |
|  | *gmmlsp1/2* | 43.10 ± 3.45* | 2.30 ± 0.67 | 31.90 ± 6.42* | 11.20 ± 3.14* | 11.81 ± 0.26* |
|  | *GmMLSP1-OE#1* | 63.30 ± 4.47* | 5.00 ± 1.05* | 77.10 ± 17.86* | 29.59 ± 7.33* | 15.92 ± 0.54* |
|  | *GmMLSP1-OE#2* | 68.90 ± 5.99* | 5.60 ± 1.84* | 83.40 ± 14.02* | 29.20 ± 6.86* | 16.90 ± 0.72* |
|  | *GmMLSP2-OE#1* | 61.00 ± 3.06* | 3.40 ± 1.08 | 68.40 ± 14.42* | 34.13 ± 6.25* | 16.85 ± 0.52* |
|  | *GmMLSP2-OE#2* | 68.30 ± 4.79* | 3.80 ± 1.48 | 68.90 ± 19.19* | 34.09 ± 7.51* | 17.33 ± 0.52* |
| Season 2 |  |  |  |  |  |  |
|  | W82 | 35.30 ± 2.00 | 3.70 ± 0.67 | 36.70 ± 5.14 | 12.13 ± 2.00 | 15.73 ± 1.09 |
|  | *gmmlsp1* | 33.60 ± 2.88 | 3.50 ± 0.85 | 35.30 ± 3.27 | 11.52 ± 1.83 | 15.38 ± 0.77 |
|  | *gmmlsp1/2* | 26.10 ± 2.38* | 2.30 ± 0.48* | 25.60 ± 2.84* | 5.88 ± 1.54* | 11.88 ± 0.70* |
|  | *GmMLSP1-OE#1* | 38.80 ± 2.74* | 4.50 ± 0.71 | 48.90 ± 9.70* | 18.06 ± 2.35* | 19.30 ± 0.43* |
|  | *GmMLSP1-OE#2* | 38.60 ± 2.76* | 3.80 ± 1.03 | 46.10 ± 6.95* | 16.01 ± 2.33* | 19.65 ± 1.35* |
|  | *GmMLSP2-OE#1* | 42.60 ± 2.50* | 4.20 ± 0.79 | 47.60 ± 6.38* | 16.79 ± 3.12* | 19.08 ± 1.16* |
|  | *GmMLSP2-OE#2* | 41.10 ± 1.20* | 4.30 ± 0.82 | 52.00 ± 10.00* | 18.51 ± 2.99* | 19.40 ± 0.55* |

**Notes:**
<sup>1</sup>, Data were collected from field experiments during the 2024–2025 winter growing season in Hainan.
<sup>2</sup>, Field data for soybean Season 1 were collected during the summer of 2024 in Zhejiang, while Season 2 data were obtained during the 2024–2025 winter in Hainan.
<sup>3</sup>, Data are presented as means ± SD (rice: n = 20; soybean: n = 10 for all traits). Significant differences ( $P < 0.05$ ; ns, not significant) based on one-way ANOVA against wild-type controls (W82 for soybean, Nip for rice) are indicated by asterisks.

In soybean, the growth of the *gmmlsp1/2* double mutant was significantly inhibited in both greenhouse and field conditions (both in Zhejiang and Hainan), displaying slower growth, reduced plant height, and fewer pods. In contrast, plants overexpressing either *GmMLSP1* or *GmMLSP2* exhibited significantly enhanced growth (Fig. 5d, S8-10, Table 1). Interestingly, the *gmmlsp1* single mutant (lacking only *GmMLSP1*) showed no significant phenotypic differences compared to the wild-type control (W82), likely due to functional redundancy.

In summary, overexpression of rice and soybean *MLSP1* orthologs significantly enhanced plant growth and seed yield. Phenotypic differences between wild-type, mutant, and overexpression lines were more pronounced in field trials than in greenhouse conditions. We hypothesize that this contrast arises from environmental stressors (e.g., heat, high light, or drought) in field settings, whereas greenhouse conditions lack such abiotic pressures. This observation aligns with the proposed role of AtMLSP1 in modulating the response of plants to adverse growth conditions.

## DISCUSSION

In this study we found inactivation of the gene encoding AtMLSP1 resulted in a similar retarded growth phenotype as other mutations affecting subunits of the respiratory chain, especially as observed previously with Cytochrome c mutants (Coronel *et al*., 2025), but overexpression resulted in altered growth and responses to environmental conditions. Overexpression not only fully complemented the decrease in Cytochrome c and complex III subunits observed with inactivation, it resulted in altered vegetative growth in terms of flowering time, flower size and seed yield, and response to high light stress in Arabidopsis. In field conditions in rice and soybean both had altered growth and seed yield, demonstrating that the findings in Arabidopsis were directly translatable to crop plants. While numerous studies have demonstrated that single-gene alterations can improve crop yield, few have explored mechanisms that stabilize mitochondrial function to enhance environmental stress adaptation (Roeder *et al*., 2025).

There are several notable features about this finding, firstly similar to numerous previous findings knocking out genes encoding respiratory function, either structural genes encoding subunits of the five respiratory complexes, or factors required for assembly resulted in retarded growth. While complementation of these genes with overexpression constructs restores growth, they did not increase seed yield. In our over-expression lines protein abundance and respiration was recovered to normal wild type levels (and respiratory capacity in isolated mitochondria), so the increase in yield does not appear to be directly related to abundance of respiratory components. Membrane potential was increased in the overexpression lines but how this impacts growth and yield is as yet unclear. Another notable feature was the altered response to high light in the overexpression lines, with transcript abundance for marker gene that respond to high light all significantly much reduced in induction in the At*MLSP1* over-expression lines. This reveals an altered signaling defense equilibrium in the overexpressing lines, and such pathways interact with a variety of hormone signaling pathways. Also in the *atmlsp1* lines AOX was not induced as is normally observed with perturbation of the respiratory chain, revealing that whatever the molecular mechanism of MLSP, its absence does not trigger the well characterized mitochondrial retrograde response inducing *AOX*.

Previous studies with Cytochrome c have revealed alterations in energy signaling, the TOR and SnRK1 pathway, alterations in hormonal signaling with ABA and GA (Racca *et al*., 2018; Racca *et al*., 2022; Canal *et al*., 2024; Coronel *et al*., 2025). The underlying molecular features resulting in enhanced growth with MLSP overexpression are unknown. A variety of assembly factors have been described for the mitochondrial respiratory chain over the last 25 years, some orthologous to system in other eukaryotic lineages, some plant specific. In particular complex III and Cytochrome c in plants are somewhat different compared to other eukaryotic mitochondrial systems. Firstly, complex III assembly pathway differs in that unique for mitochondrial complex III (cytochrome bc_1_) it still uses a Twin-Arginine-Translocase (TAT) based assembly pathway (Schäfer *et al*., 2020), that is utilized in bacterial cytochrome bc_1_ and chloroplast cytochrome b_6_f complex (Molik *et al*., 2001), but not in other eukaryotes (Rosales-Hernandez *et al*., 2025). Also, notably the mitochondrial processing peptidase is incorporated into complex III in plants, rather than a soluble matrix protein as observed in other eukaryotic systems (Braun *et al*., 1992; Glaser *et al*., 1994). For Cytochrome c the insertion of heme is via a type I system similar to the α-proteobacterial ancestor of mitochondria (and also for c1 in complex III) with Oxidase assembly protein 2 required for assembly (Kolli *et al*., 2020). This suggests that there may be more unique molecular components required for the assembly, disassembly and stability of complex III and Cytochrome c in plants compared to other eukaryotic systems.

Overall, what these findings reveal that alterations in respiration is a feasible approach to increase both plant productivity and environmental tolerance simultaneously. This will likely be achieved by targeting factors that are not the direct catalytic enzymes involved in respiration, but rather components associated with assembly, disassembly or stability of respiratory complexes. In this way it appears that overexpression of such factors appears to avoid stress signaling and may make the respiratory complexes more robust under unfavorable conditions. It has also been noted that altering proteolysis improves both photosynthesis and restores growth of a mutant with complex I deficiency (not completely knocked out) (Grimmer *et al*., 2020; Ivanova *et al*., 2021). Thus, stabilization of proteins to prevent degradations appears to be a viable route to increase stress tolerance and productivity.

## ACKNOWLEDGMENTS

This work was supported by the Ministry of Science and Technology of China (2023ZD04072, 2021YFF1001204, 2021YFF1000402), the National Natural Science Foundation of China (32572127), School of Life Sciences Henan University and the 111 project of the Ministry of Education (B14027). The authors would like to thank Biogle Genetech (Hangzhou Biogle Co., LTD.) for rice transformation. We especially thank Dr. Shan Zhang (School of Medicine, Zhejiang University) for insightful discussions and suggestions. We also thank Drs. Yong Wang, Shelong Zhang, Jiming Xu, Zhenyu Qi and Rui Sun for technical assistance and field experiments in the Agricultural Experiment Stations of Zhejiang university. We thank Prof. Xueqin Zhang from China Agricultural University for supporting Arabidopsis germplasm used in this study.

## COMPETING INTERESTS

The authors declare no competing interests.

## AUTHOR CONTRIBUTIONS

J.W., H.S. and L.G. conceived the project. L.G. performed experiments with Y.Z. contributing with respiratory measurements. L.G., Y.Z., X.W., J.W. and H.S. interpreted results and drafted the publication. All authors reviewed the manuscript.

## DATA AVAILABILITY

Sequence data from this article can be found in the TAIR or Phytozome database under the following accession numbers: *AtMLSP1 (At4g17085)*, *OsMLSP1 (LOC_Os06g50090)*, *GmMLSP1 (Glyma.05g067900)*, *GmMLSP2 (Glyma.17g150000)*, *NDUFS4 (At5g67590)*, *COXII (AtMg00160)*, *ATP β (At5g08670)*, *TOM40 (At3g20000)*, *Tom20-3 (At3g27080)*, *RISP (At5g13430)*, *TIM23-1 (At1g17530)*, *Cyt c-1 (At1g22840)*, *Cyt c-2 (At4g10040)*. *ELIP1 (AT3G22840)*, *ELIP2 (AT4G14690)*, *HSP17.6 (AT5G12030)*, *PAP1 (AT1G56650)*.

## SUPPORTING INFORMATION

**Figure S1.** AtMLSP1 is a mitochondrial Protein. Public published studies showing a mitochondrial location of AtMLSP1.

**Figure S2.** Rosette leaves number of Col-0, mutants and overexpressing plants grown on soil and phenotypic analysis of plants grown on plates in a 14-h-light/10-h-dark cycle for germination and early seedling growth. Growth stage was performed according to Boyes et al., 2002 (Boyes *et al*., 2001).

**Figure S3.** Generation of mutation lines for *Arabidopsis thaliana*.

**Figure S4.** Generation of mutation lines for *Glycine Max* (Soybean).

**Figure S5.** Generation of mutation lines for *Oryza sativa* (Rice).

**Figure S6.** Knock out of *OsMLSP1* inhibited growth of rice in green house.

**Figure S7.** *MLSP1*-overexpressing transgenic rice exhibits improved yield in field.

**Figure S8.** Phenotype of soybean materials in green house.

**Figure S9.** *MLSP* overexpressing transgenic soybean exhibits improved yield in field (Zhejiang).

**Figure S10.** *MLSP* overexpressing transgenic soybean exhibits improved yield in field (Hainan).

**Table S1.** List of antibodies used in this study.

**Table S2.** Primers used in this study.

## Notes

### Competing Interest Statement

The authors have declared no competing interest.

