## Supplemental Figure S1 for "A NOVEL MITOCHONDRIAL PEPTIDE ESSENTIAL FOR RESPIRATORY CAPACITY PROMOTES GROWTH, YIELD AND ABIOTIC STRESS TOLERANCE IN PLANTS"

(a)

| Published research about mitochondria proteomics | Experimental Localizations (MS/MS) |
| --- | --- |
| Tan et al., 2012 | Mitochondrion |
| Senkler et al., 2017 | Mitochondrion |
| Rugen et al., 2019 | Mitochondrion |
| Kuhnert et al., 2020 | Mitochondrion |
| Fuchs et al., 2020 | Mitochondrion |

(b)

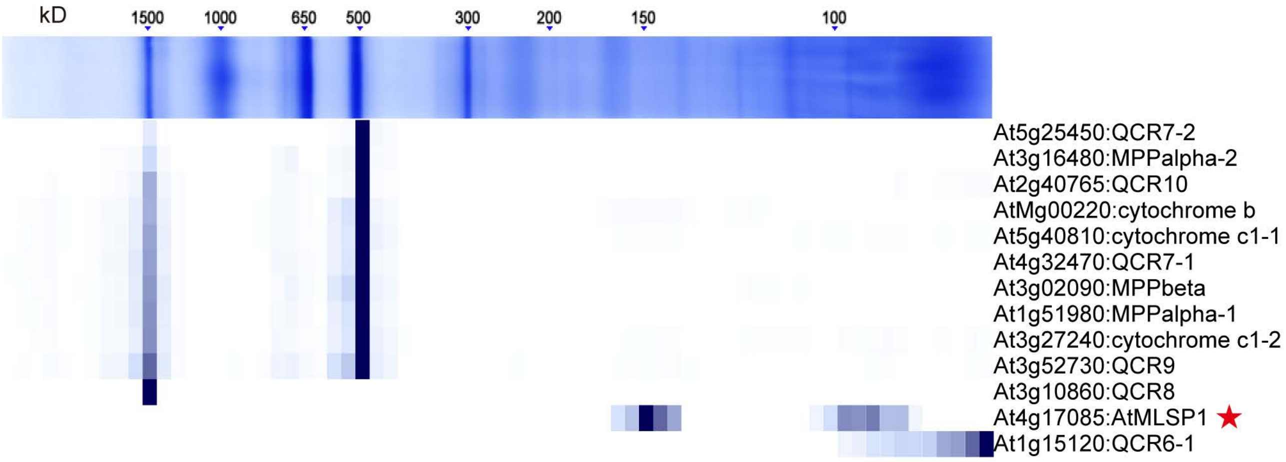


(c)

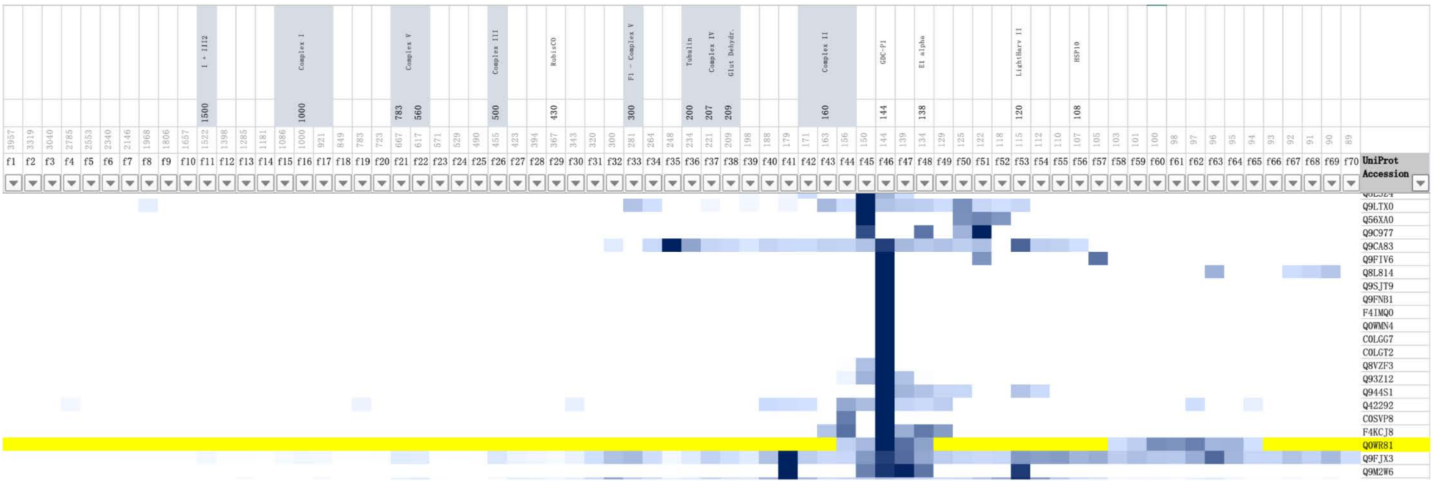

**Figure S1.** AtMLSP1 is a mitochondrial Protein. Public published studies showing a mitochondrial location of AtMLSP1.

**(a)** Published studies on the mitochondrial proteome from Arabidopsis identifying AtMLSP1 as a mitochondrial protein based on mass spectrometry identification of mitochondrial proteins. **(b-c)** GelMap images showing the identification of AtMLSP1 as a mitochondrial protein.
