## Supplemental Figure S2 for "A NOVEL MITOCHONDRIAL PEPTIDE ESSENTIAL FOR RESPIRATORY CAPACITY PROMOTES GROWTH, YIELD AND ABIOTIC STRESS TOLERANCE IN PLANTS"

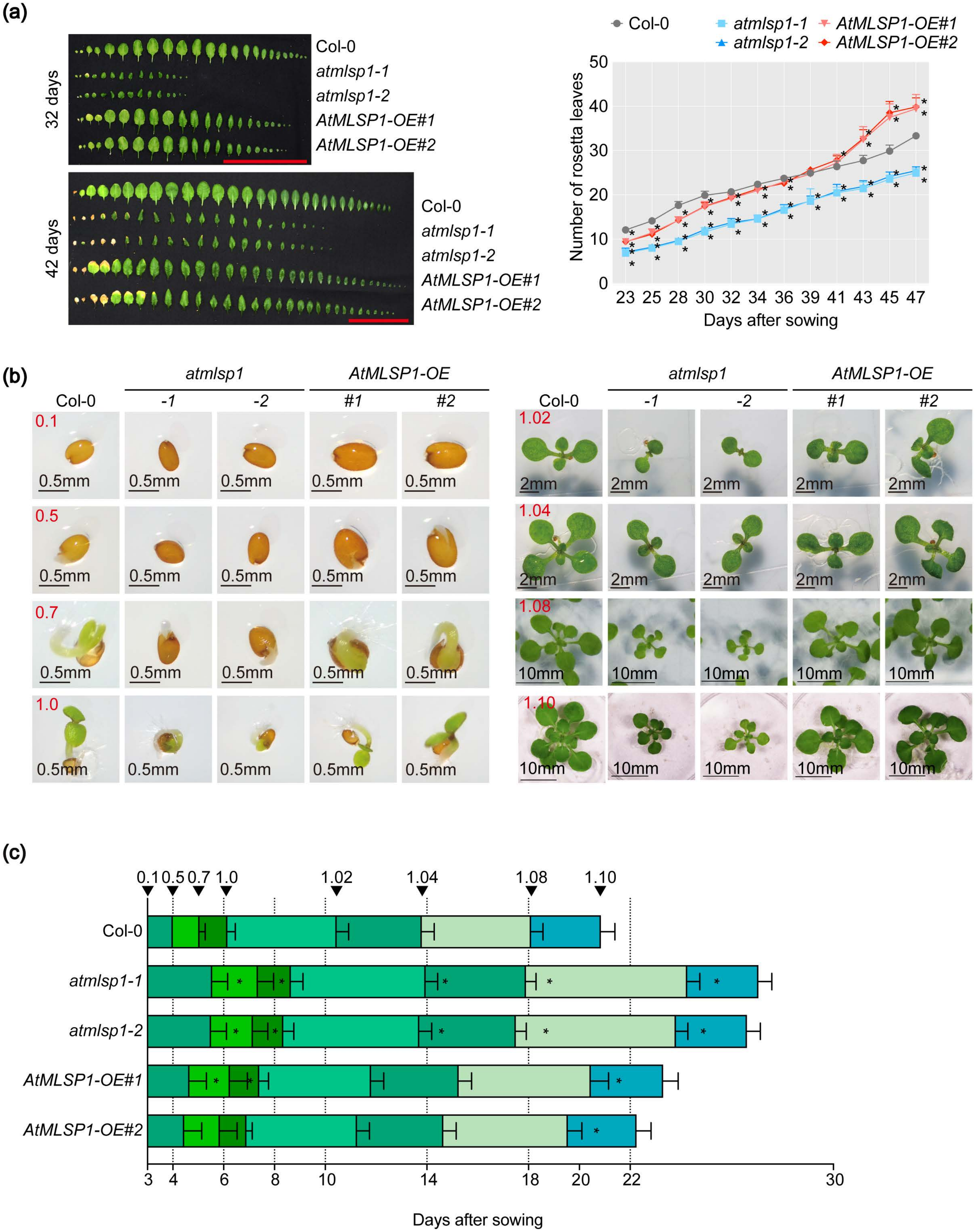

(a)
32 days
42 days
Col-0
atmlsp1-1
atmlsp1-2
AtMLSP1-OE#1
AtMLSP1-OE#2
Number of rosetta leaves
Days after sowing
(b)
atmlsp1
AtMLSP1-OE
Col-0
-1
-2
#1
#2
0.1
0.5
0.7
1.0
1.02
1.04
1.08
1.10
0.5mm
2mm
10mm
(c)
Days after sowing

**Figure S2.** Rosette leaves number of Col-0, mutants and overexpressing plants grown on soil and phenotypic analysis of plants grown on plates in a 14-h-light/10-h-dark cycle for germination and early seedling growth**.** Growth stage was performed according to Boyes et al., 2002 (Boyes *et al.*, 2001).

**(a)** Leaf numbers and phenotypes for analysed genotypes. Scale bar, 10 cm. Values are means ± SDs (n = 20; one-way ANOVA; *P < 0.05). **(b-c)** Representative images of growth stage analysis up to stage 1.10 for the Arabidopsis lines. Scale bar, as indicated on the images. Figure below showed growth stage progression in the analysed genotypes. Values are means ± SDs (n = 20; one-way ANOVA; *P < 0.05).
