## Supplemental Figure S3 for "A NOVEL MITOCHONDRIAL PEPTIDE ESSENTIAL FOR RESPIRATORY CAPACITY PROMOTES GROWTH, YIELD AND ABIOTIC STRESS TOLERANCE IN PLANTS"

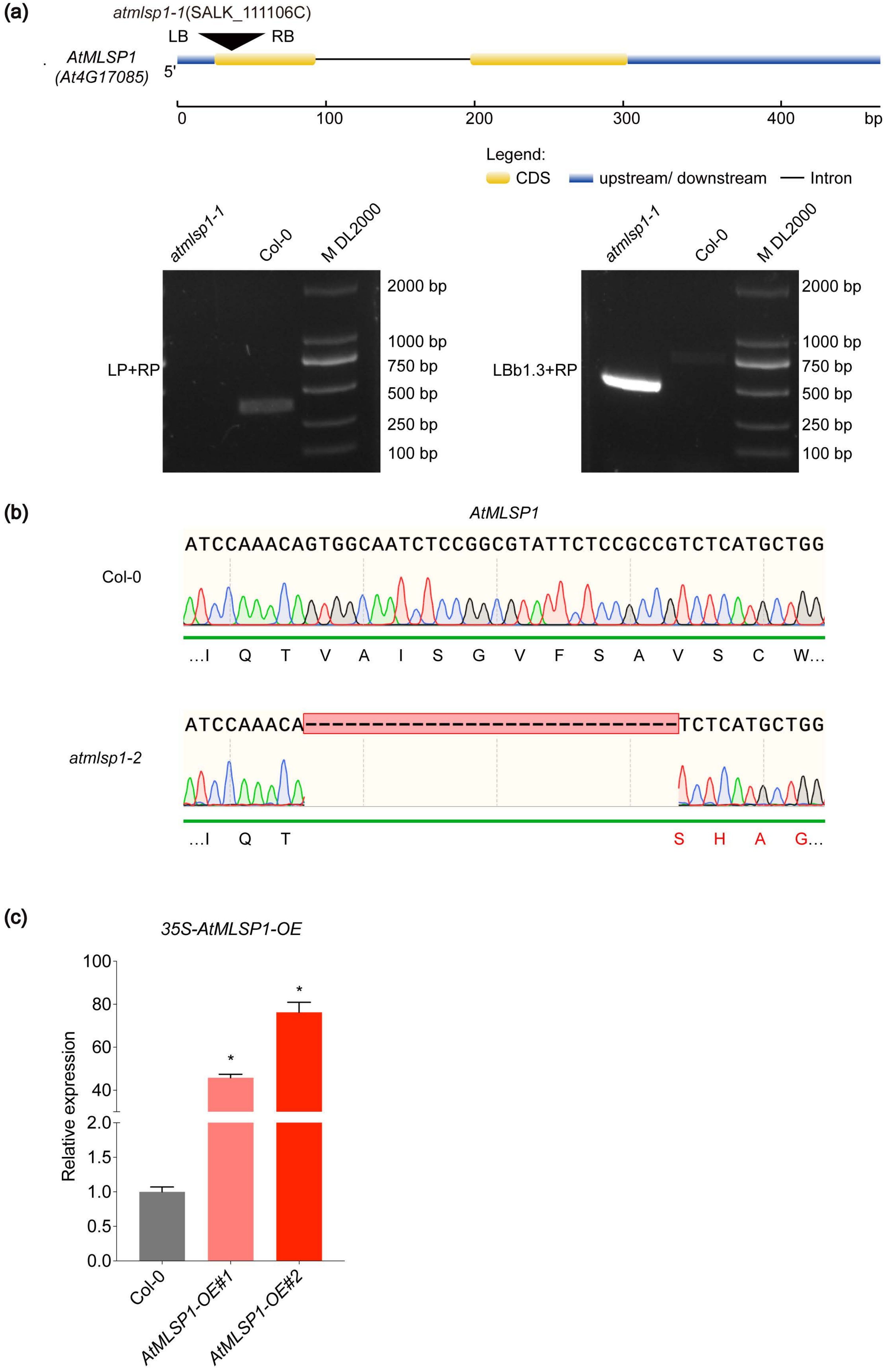
(a)
atmlsp1-1(SALK_111106C)
LB
RB
AtMLSP1
(At4G17085)
5'
0
100
200
300
400
bp
Legend:
CDS
upstream/ downstream
Intron
atmlsp1-1
Col-0
M DL2000
LP+RP
2000 bp
1000 bp
750 bp
500 bp
250 bp
100 bp
atmlsp1-1
Col-0
M DL2000
LBb1.3+RP
2000 bp
1000 bp
750 bp
500 bp
250 bp
100 bp
(b)
AtMLSP1
Col-0
ATCCAAACAGTGGCAATCTCCGGCGTATTCTCCGCCGTCTCATGCTGG
…I Q T V A I S G V F S A V S C W…
atmlsp1-2
ATCCAAACA-----------------------------------TCTCATGCTGG
…I Q T S H A G…
(c)
35S-AtMLSP1-OE
Relative expression
100
80
60
40
2.0
1.5
1.0
0.5
0.0
*
*
Col-0
AtMLSP1-OE#1
AtMLSP1-OE#2

**Figure S3.** Generation of mutation lines for *Arabidopsis thaliana*.

**(a)** The schematic gene model of *AtMLSP1* showing the T-DNA insertion site in *atmlsp1-1* and identification of the *atmlsp1-1* homozygous mutant by PCR. LP and RP are primers flanking *AtMLSP1*, and LBb1.3 is the T-DNA-specific primer. **(b)** Sanger sequencer analysis of Col-0 and *atmlsp1-2*, which confirmed the 28 bp deletion that caused amino acid changes. **(c)** qRT-PCR verification of *AtMLSP1* transcript levels in leaves of CaMV35S promoter-driven overexpression lines. Data were expressed as means ± SD from three biologicals and two technical replicates. Significant difference (*$P < 0.05$) based on one-way ANOVA compared with Col-0 are indicated by asterisks.
