## Supplemental Figure S4 for "A NOVEL MITOCHONDRIAL PEPTIDE ESSENTIAL FOR RESPIRATORY CAPACITY PROMOTES GROWTH, YIELD AND ABIOTIC STRESS TOLERANCE IN PLANTS"

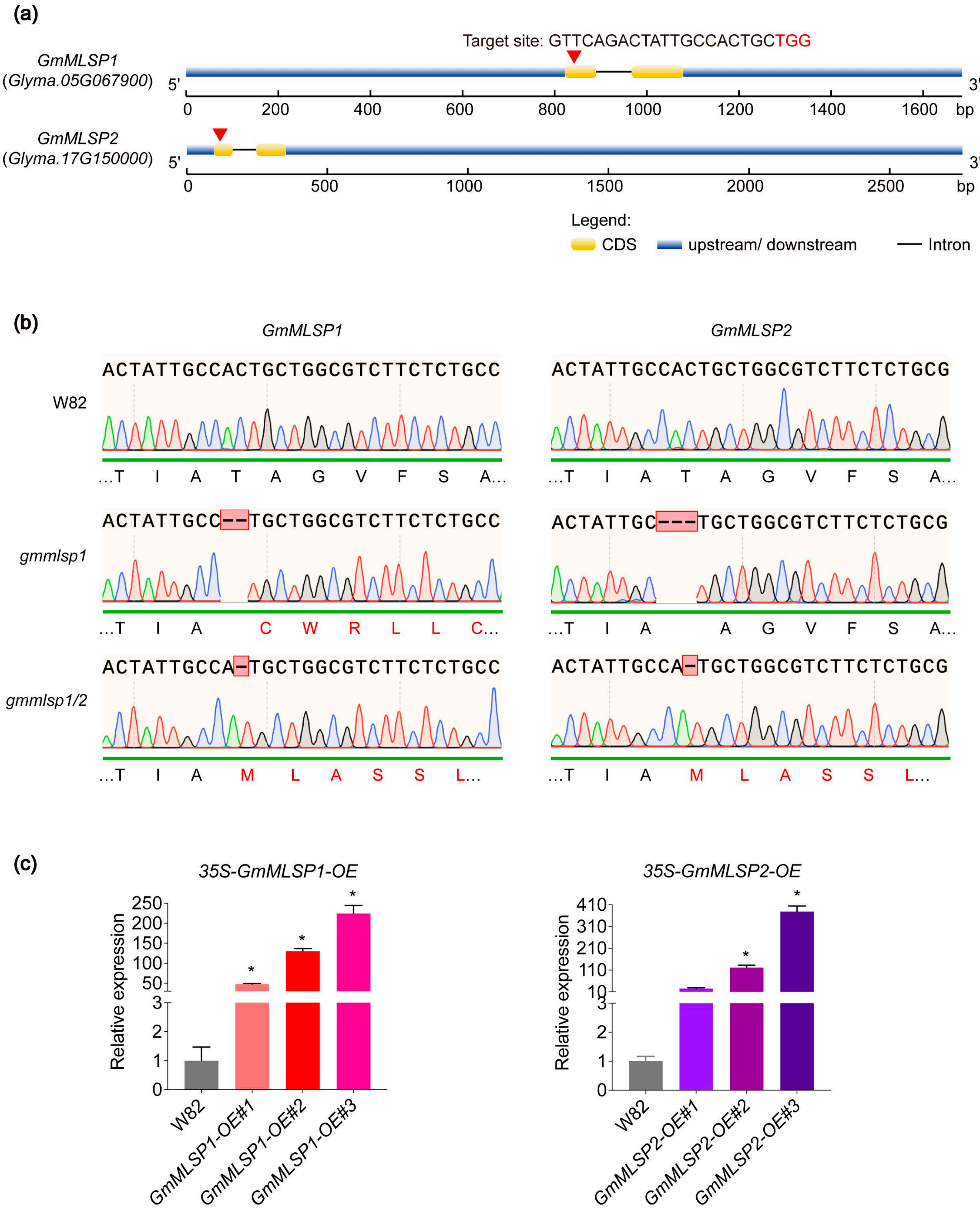

(a)
Target site: GTTCAGACTATTGCCACTGCTGG
GmMLSP1
(Glyma.05G067900)
5'
3'
0
200
400
600
800
1000
1200
1400
1600
bp
GmMLSP2
(Glyma.17G150000)
0
500
1000
1500
2000
2500
Legend:
CDS
upstream/ downstream
Intron
(b)
GmMLSP1
GmMLSP2
W82
ACTATTGCCACTGCTGGCGTCTTCTCTGCC
ACTATTGCCACTGCTGGCGTCTTCTCTGCG
...T I A T A G V F S A...
gmmlsp1
ACTATTGCC--TGCTGGCGTCTTCTCTGCC
...T I A C W R L L C...
ACTATTGC---TGCTGGCGTCTTCTCTGCG
...T I A A G V F S A...
gmmlsp1/2
ACTATTGCCA-TGCTGGCGTCTTCTCTGCC
ACTATTGCCA-TGCTGGCGTCTTCTCTGCG
...T I A M L A S S L...
(c)
35S-GmMLSP1-OE
35S-GmMLSP2-OE
Relative expression
250
200
150
100
50
3
2
1
0
410
310
210
110
10
*
W82
GmMLSP1-OE#1
GmMLSP1-OE#2
GmMLSP1-OE#3
GmMLSP2-OE#1
GmMLSP2-OE#2
GmMLSP2-OE#3

**Figure S4.** Generation of mutation lines for *Glycine Max* (Soybean).

**(a)** Schematic gene models of *GmMLSP1* and *GmMLSP2* with sequence for generation of CRISPR-Cas9 mutants shown in figure (PAM sequence is highlighted in red color). **(b)** Sanger sequencer analysis of W82, *gmmlsp1* and *gmmlsp1/2,* which confirmed the mutations that caused amino acid changes. **(c)** qRT-PCR verification of *GmMLSP1* or *GmMLSP2* transcript levels in leaves of CaMV35S promoter-driven overexpression lines. Data were expressed as means ± SD from three biologicals and two technical replicates. Significant difference (*$P < 0.05$) based on one-way ANOVA compared with W82 are indicated by asterisks.
