## Supplemental Figure S5 for "A NOVEL MITOCHONDRIAL PEPTIDE ESSENTIAL FOR RESPIRATORY CAPACITY PROMOTES GROWTH, YIELD AND ABIOTIC STRESS TOLERANCE IN PLANTS"

(a)

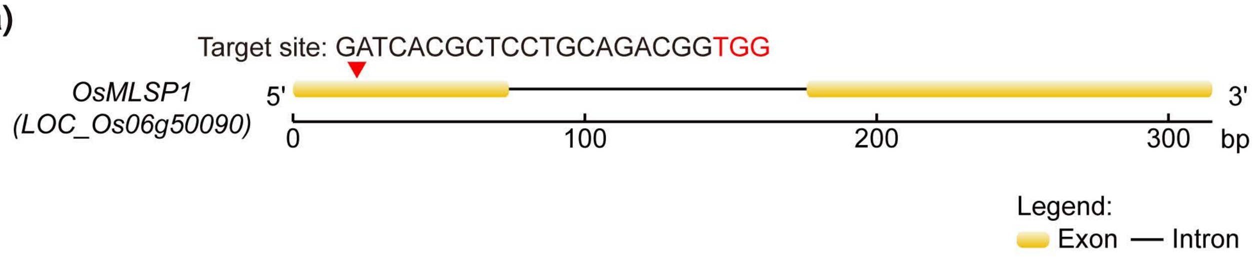

Target site: GATCACGCTCCTGCAGACGGTGG
OsMLSP1
(LOC_Os06g50090)
5'
3'
0
100
200
300
bp
Legend:
Exon
Intron


(b)

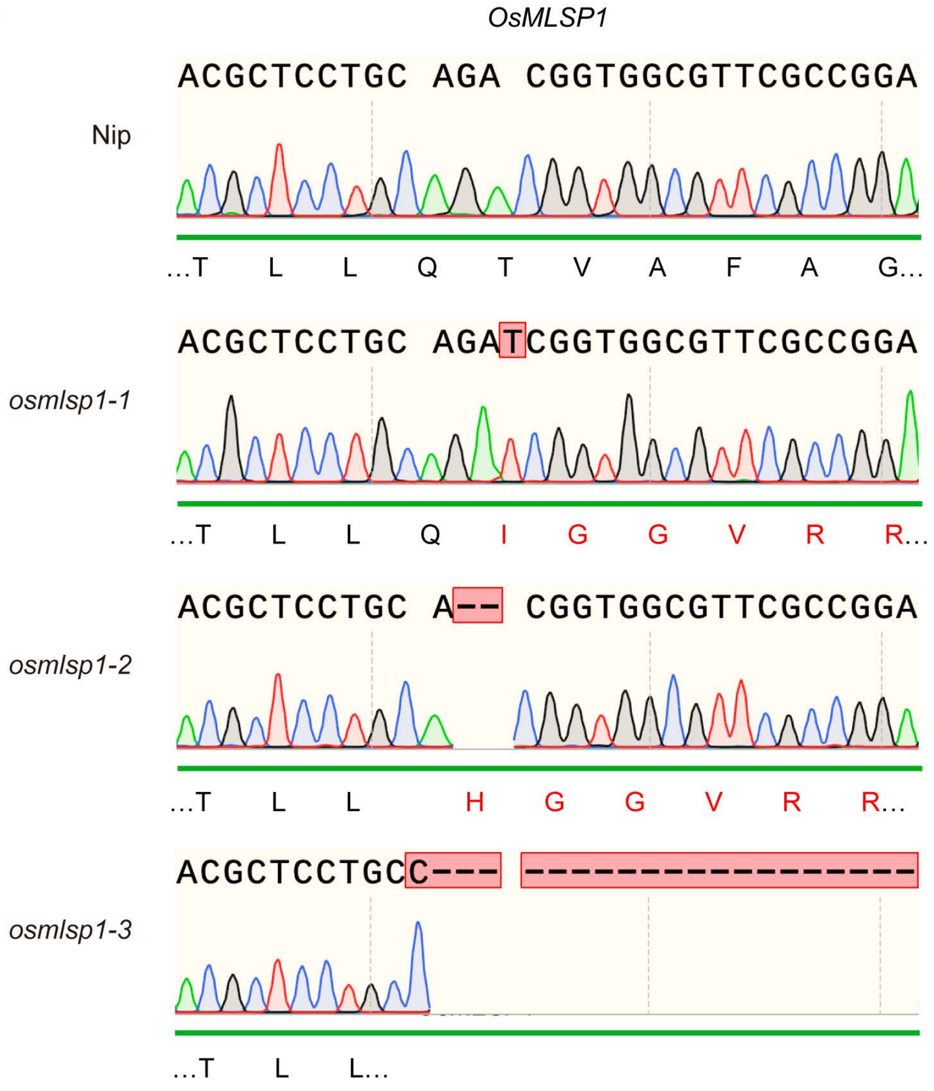

OsMLSP1
Nip
ACGCTCCTGC AGA CGGTGGCGTTCGCCGGA
...T L L Q T V A F A G...
osmlsp1-1
ACGCTCCTGC AGATCGGTGGCGTTCGCCGGA
...T L L Q I G G V R R...
osmlsp1-2
ACGCTCCTGC A-- CGGTGGCGTTCGCCGGA
...T L L H G G V R R...
osmlsp1-3
ACGCTCCTGCC--- -----------------
...T L L...


(c)

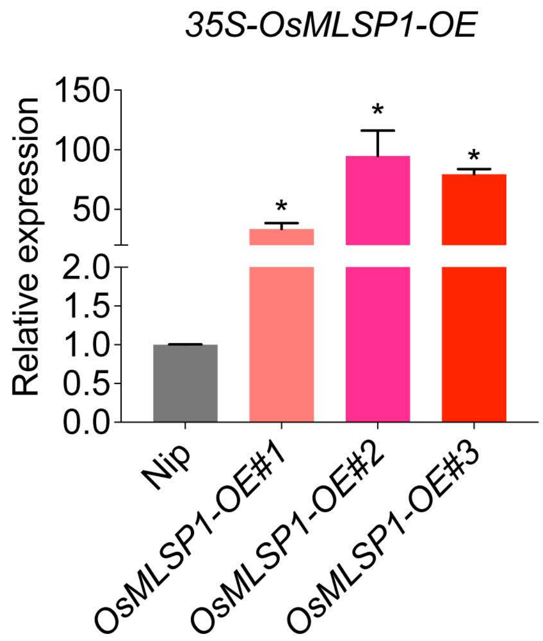

35S-OsMLSP1-OE
Relative expression
150
100
50
2.0
1.5
1.0
0.5
0.0
*
Nip
OsMLSP1-OE#1
OsMLSP1-OE#2
OsMLSP1-OE#3

**Figure S5.** Generation of mutation lines for *Oryza sativa* (Rice).

**(a)** Schematic gene models of *OsMLSP1* with sequence for generation of CRISPR-Cas9 mutants shown in figure (PAM sequence is highlighted in red color). **(b)** Sanger sequencer analysis of Nip, *osmlsp1-1*, *osmlsp1-2* and *osmlsp1-3*, which confirmed the mutations that caused amino acid changes. **(c)** qRT-PCR verification of *OsMLSP1* transcript levels in leaves of CaMV35S promoter-driven overexpression lines. Data were expressed as means ± SD from three biologicals and two technical replicates. Significant difference (*P < 0.05) based on one-way ANOVA compared with Nip are indicated by asterisks.
