## Supplemental Figure S6 for "A NOVEL MITOCHONDRIAL PEPTIDE ESSENTIAL FOR RESPIRATORY CAPACITY PROMOTES GROWTH, YIELD AND ABIOTIC STRESS TOLERANCE IN PLANTS"

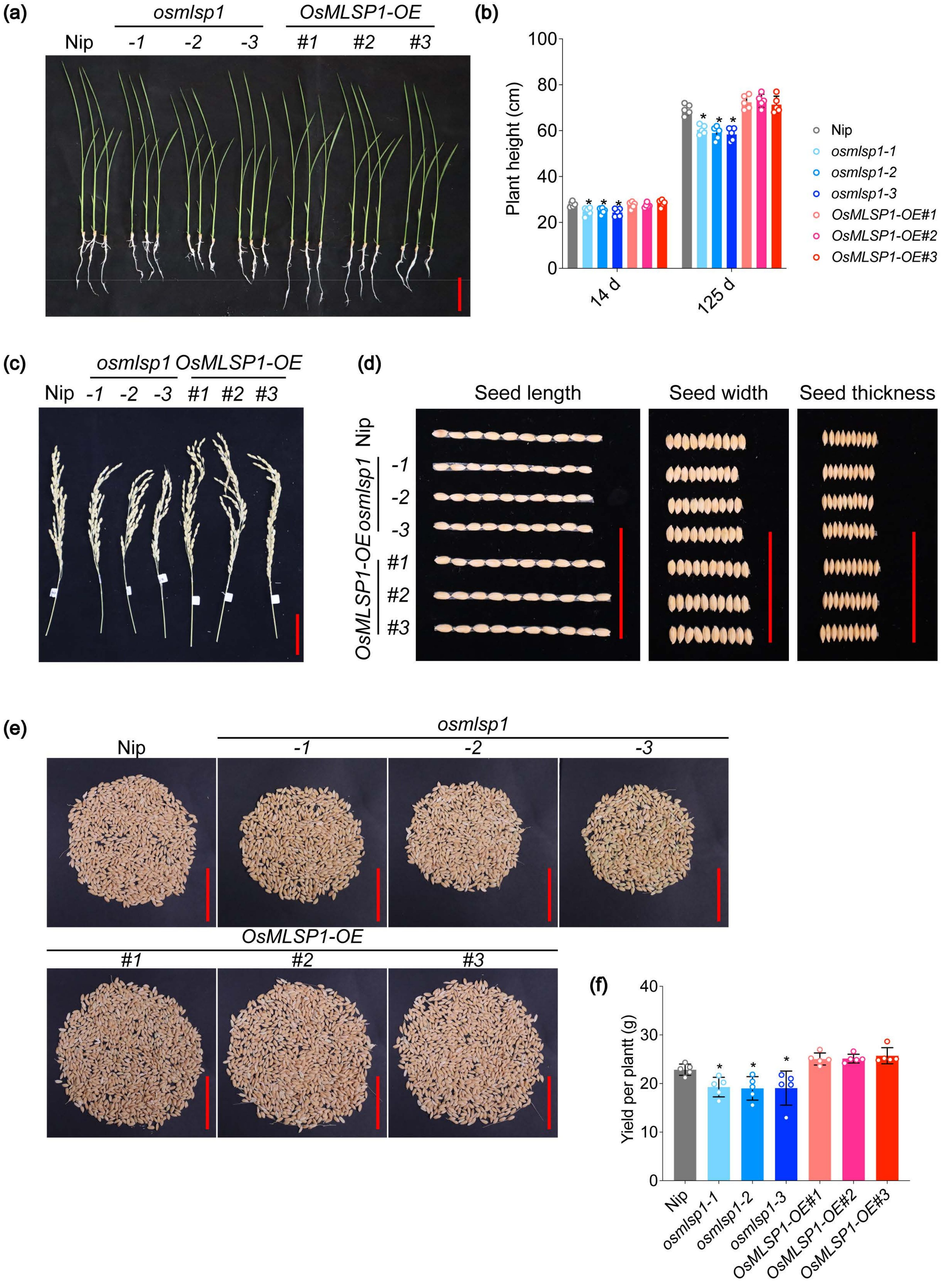

(a)
osmlsp1
OsMLSP1-OE
Nip
-1
-2
-3
#1
#2
#3
(b)
Plant height (cm)
100
80
60
40
20
0
14 d
125 d
Nip
osmlsp1-1
osmlsp1-2
osmlsp1-3
OsMLSP1-OE#1
OsMLSP1-OE#2
OsMLSP1-OE#3
(c)
osmlsp1 OsMLSP1-OE
Nip -1 -2 -3 #1 #2 #3
(d)
Seed length
Seed width
Seed thickness
Nip
osmlsp1
OsMLSP1-OE
(e)
Nip
osmlsp1
-1
-2
-3
OsMLSP1-OE
#1
#2
#3
(f)
Yield per plantt (g)
40
30
20
10
0
Nip
osmlsp1-1
osmlsp1-2
osmlsp1-3
OsMLSP1-OE#1
OsMLSP1-OE#2
OsMLSP1-OE#3

**Figure S6.** Knock out of *OsMLSP1* inhibited growth of rice in green house.

**(a)** Growth of hydroponic cultured Nip, mutants and *MLSP1*-overexpressing rice at 14 d after emergency. Scale bar, 5 cm. **(b)** Plant height of hydroponic cultured or Nip, mutants and *MLSP1*-overexpressing rice. Different letters indicate significant differences (n = 5; one-way ANOVA; *P < 0.05). **(c)** Primary panicle morphology of Nip, mutants and *MLSP1*-overexpressing rice in green house. Scale bar, 5 cm. **(d)** Seed morphology of Nip, mutants and *MLSP1*-overexpressing rice in green house. Scale bar, 5 cm. **(e-f)** Single-plant yield of Nip, mutants and *MLSP1*-overexpressing rice in green house. Scale bar, 5 cm. Different letters indicate significant differences (n = 5; one-way ANOVA; *P < 0.05).
