## Supplemental Figure S7 for "A NOVEL MITOCHONDRIAL PEPTIDE ESSENTIAL FOR RESPIRATORY CAPACITY PROMOTES GROWTH, YIELD AND ABIOTIC STRESS TOLERANCE IN PLANTS"

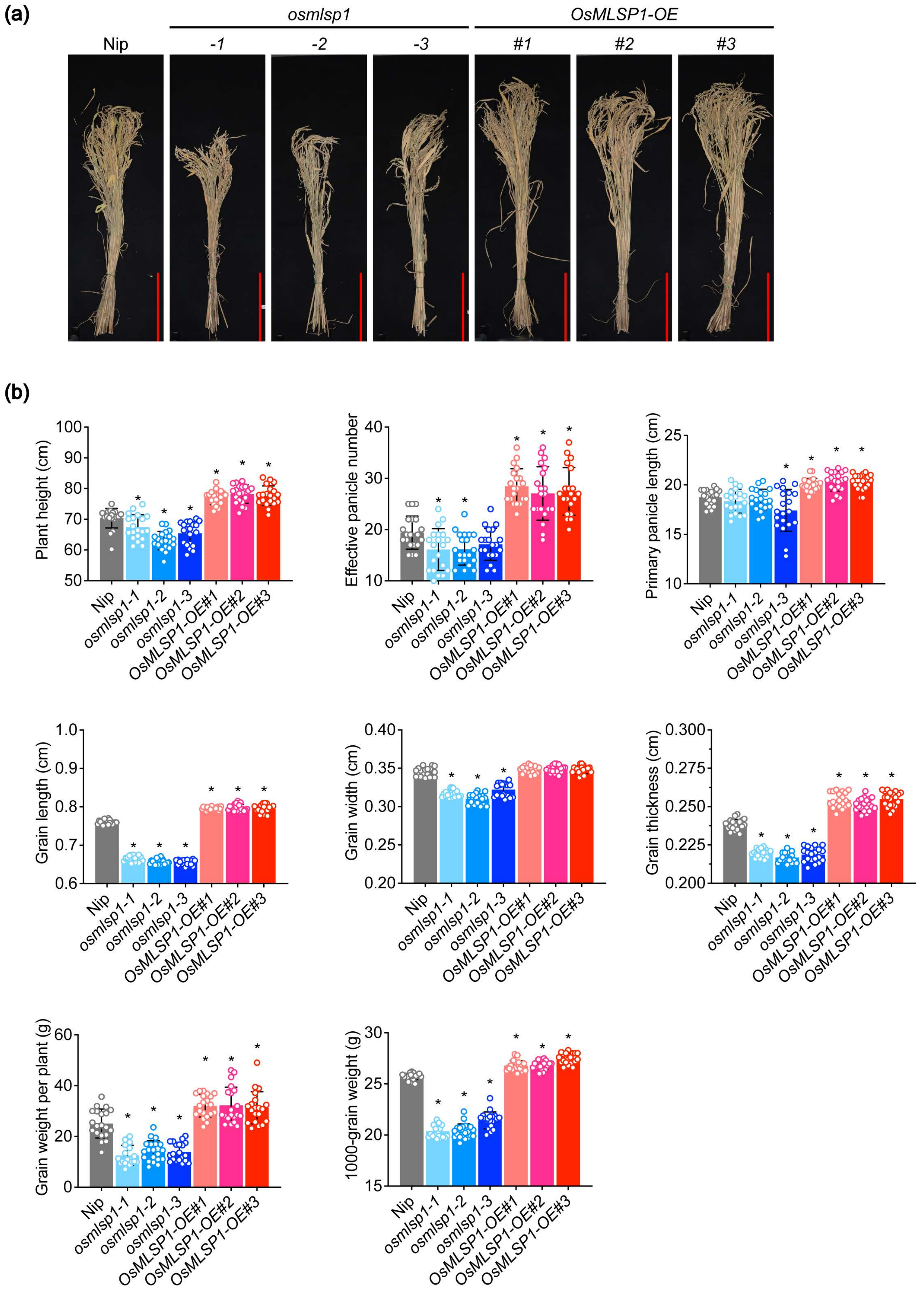

(a)
osmlsp1
OsMLSP1-OE
Nip
-1
-2
-3
#1
#2
#3
(b)
Plant height (cm)
Effective panicle number
Primary panicle length (cm)
Grain length (cm)
Grain width (cm)
Grain thickness (cm)
Grain weight per plant (g)
1000-grain weight (g)
Nip
osmlsp1-1
osmlsp1-2
osmlsp1-3
OsMLSP1-OE#1
OsMLSP1-OE#2
OsMLSP1-OE#3

**Figure S7.** *MLSP1*-overexpressing transgenic rice exhibits improved yield in field.

**(a)** Growth of Nip, mutants and *MLSP1*-overexpressing rice in Hainan after harvested. Scale bar, 20 cm. **(b)** Statistical analysis of plant height, effective panicle number per plant, primary panicle length, grain yield per plant, 1000-grain weight, grain length, grain width, grain thickness of Nip, mutants and *MLSP1*-overexpressing rice in Hainan. Values are means ± SDs (n = 20; one-way ANOVA; *P < 0.05).
