## Supplemental Figure S8 for "A NOVEL MITOCHONDRIAL PEPTIDE ESSENTIAL FOR RESPIRATORY CAPACITY PROMOTES GROWTH, YIELD AND ABIOTIC STRESS TOLERANCE IN PLANTS"

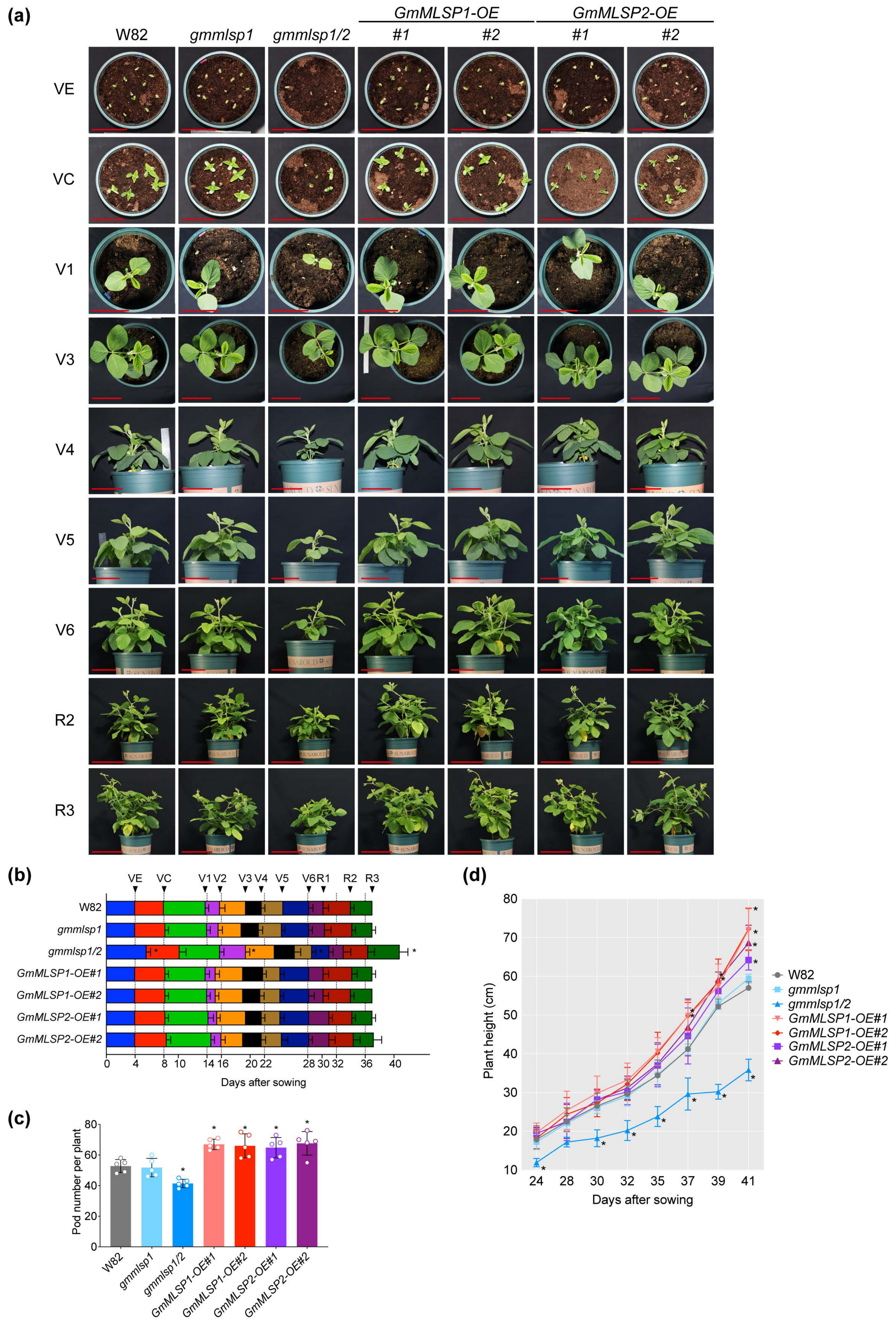
(a)
GmMLSP1-OE
GmMLSP2-OE
W82
gmmlsp1
gmmlsp1/2
#1
#2
#1
#2
VE
VC
V1
V3
V4
V5
V6
R2
R3
(b)
VE VC V1 V2 V3 V4 V5 V6 R1 R2 R3
W82
gmmlsp1
gmmlsp1/2
GmMLSP1-OE#1
GmMLSP1-OE#2
GmMLSP2-OE#1
GmMLSP2-OE#2
0 4 8 10 14 16 20 22 28 30 32 36 40
Days after sowing
(c)
Pod number per plant
0 20 40 60 80
W82
gmmlsp1
gmmlsp1/2
GmMLSP1-OE#1
GmMLSP1-OE#2
GmMLSP2-OE#1
GmMLSP2-OE#2
(d)
Plant height (cm)
10 20 30 40 50 60 70 80
24 28 30 32 35 37 39 41
Days after sowing
W82
gmmlsp1
gmmlsp1/2
GmMLSP1-OE#1
GmMLSP1-OE#2
GmMLSP2-OE#1
GmMLSP2-OE#2

**Figure S8.** Phenotype of soybean materials in green house.

**(a)** Vegetative and floral development of W82, mutants and *MLSPs*-overexpressing soybean plants under greenhouse conditions. VC (vegetative stage, cotyledon), V1-V6 (vegetative stage, first to sixth node). Scale bar, 11.5 cm. R2 (reproductive stage, full bloom) and R3 (reproductive stage, beginning pod). Scale bar, 23 cm. **(b)** Growth stage progression in the analysed genotypes. 5 plants were selected for analysis. VE-R3 as mentioned above. Values are means ± SDs (n = 5; one-way ANOVA; $*P < 0.05$). **(c)** Pod number of W82, mutants and *MLSPs*-overexpressing soybean plants in green house. Values are means ± SDs (n = 5; one-way ANOVA; $*P < 0.05$). **(d)** Seed morphology of W82, mutants and *MLSPs*-overexpressing soybean plants. Scale bar, 5 cm.
