## Supplemental Figure S9 for "A NOVEL MITOCHONDRIAL PEPTIDE ESSENTIAL FOR RESPIRATORY CAPACITY PROMOTES GROWTH, YIELD AND ABIOTIC STRESS TOLERANCE IN PLANTS"

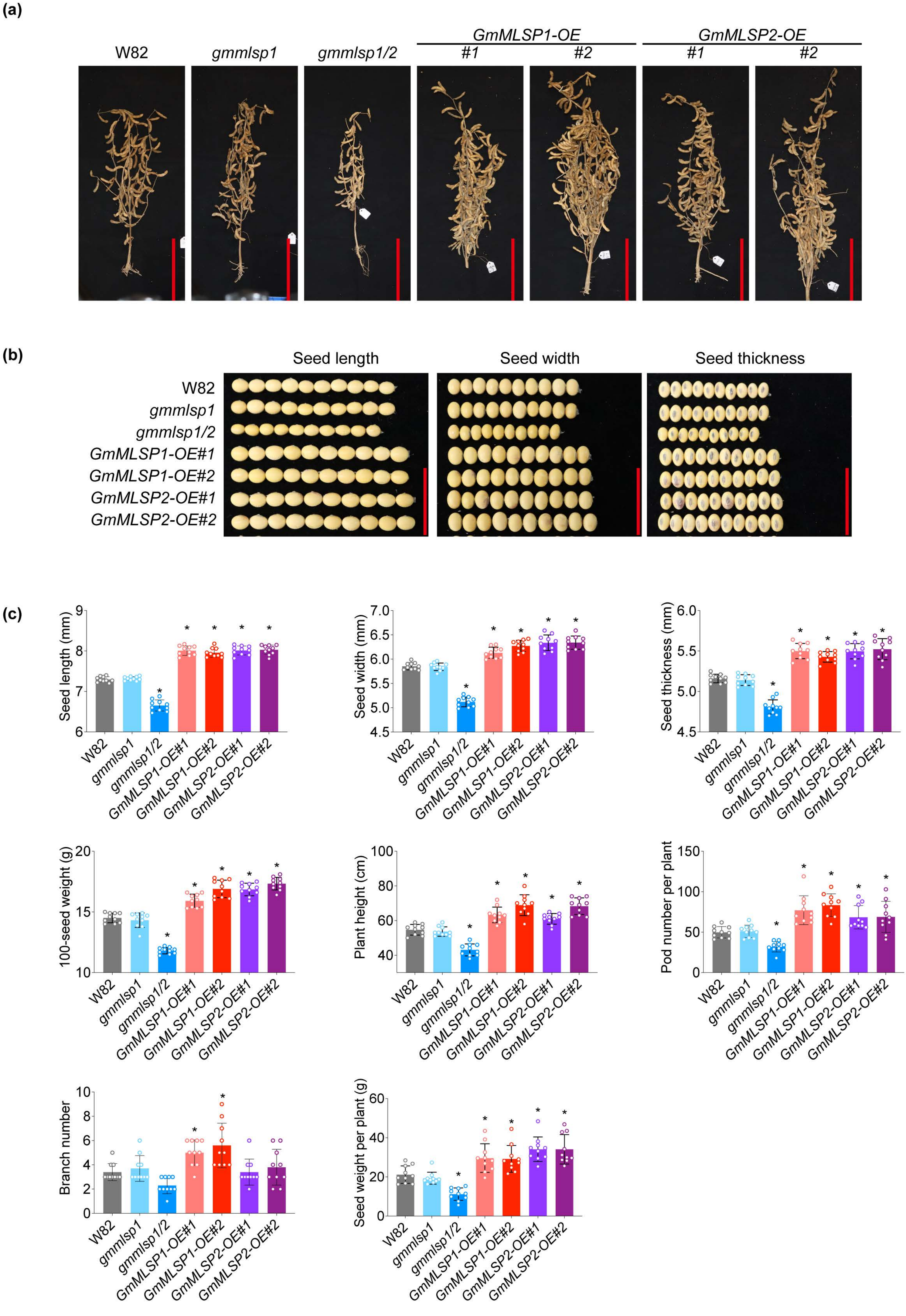
(a)
W82
gmmlsp1
gmmlsp1/2
GmMLSP1-OE
#1
#2
GmMLSP2-OE
#1
#2
(b)
Seed length
Seed width
Seed thickness
W82
gmmlsp1
gmmlsp1/2
GmMLSP1-OE#1
GmMLSP1-OE#2
GmMLSP2-OE#1
GmMLSP2-OE#2
(c)
Seed length (mm)
Seed width (mm)
Seed thickness (mm)
100-seed weight (g)
Plant height (cm)
Pod number per plant
Branch number
Seed weight per plant (g)
W82
gmmlsp1
gmmlsp1/2
GmMLSP1-OE#1
GmMLSP1-OE#2
GmMLSP2-OE#1
GmMLSP2-OE#2

**Figure S9.** *MLSP* overexpressing transgenic soybean exhibits improved yield in field (Zhejiang).

**(a)** Growth of W82, mutants and *MLSPs*-overexpressing soybean plants in Zhejiang after harvested. Scale bar, 20 cm. **(b)** Seed morphology of W82, mutants and *MLSPs*-overexpressing soybean plants in Zhejiang. Scale bar, 3 cm. **(c)** Statistical analysis of seed length, seed width, seed thickness, 100-seed weight, plant height, pod number per plant, branch number and seed weight per plant of W82, mutants and *MLSPs*-overexpressing soybean plants in Zhejiang. Values are means ± SDs ($n = 10$; one-way ANOVA; $^*P < 0.05$).
